# The Drosophila MIC10 orthologue has a propensity to polymerize into cristae-shaping filaments

**DOI:** 10.1101/2023.04.17.537183

**Authors:** Till Stephan, Stefan Stoldt, Mariam Barbot, Travis D. Carney, Felix Lange, Mark Bates, Peter Bou Dib, Halyna R. Shcherbata, Michael Meinecke, Dietmar Riedel, Sven Dennerlein, Peter Rehling, Stefan Jakobs

**Affiliations:** Department of NanoBiophotonics, Max Planck Institute for Multidisciplinary Sciences, Göttingen, Germany; Clinic of Neurology, University Medical Center Göttingen, Göttingen, Germany; Cluster of Excellence “Multiscale Bioimaging: from Molecular Machines to Networks of Excitable Cells” (MBExC), University of Göttingen, Germany; Institute of Cell Biochemistry, Hannover Medical School, Hanover, Germany; Mount Desert Island Biological Laboratory, Bar Harbor, ME, United States; Department of Optical Nanoscopy, Institute for Nanophotonics, Göttingen, Germany; Biochemistry Center (BZH), Heidelberg University, Heidelberg, Germany; Laboratory of Electron Microscopy, Max Planck Institute for Multidisciplinary Science, Göttingen, Germany; Department of Cellular Biochemistry, University Medical Center Göttingen, Göttingen, Germany; Fraunhofer Institute for Translational Medicine and Pharmacology, Translational Neuroinflammation and Automated Microscopy, Göttingen, Germany; Max Planck Institute for Multidisciplinary Science, 37077 Göttingen, Germany

## Abstract

Mitochondria are essential eukaryotic double-membrane organelles. The convoluted mitochondrial inner membrane forms highly organized invaginations, termed cristae, which are crucial for energy metabolism. Cristae formation requires MICOS, a conserved hetero-oligomeric inner membrane complex. The MICOS core subunit MIC10 is a small transmembrane protein that oligomerizes through highly conserved glycine-rich motifs to control cristae formation. Sequence alignments show that *D. melanogaster* exhibits three MIC10-like proteins with different tissue-specific expression patterns. Here, we show that the ubiquitously expressed Dmel_CG41128/MINOS1b/DmMIC10b is the major MIC10 orthologue in flies. Loss of DmMIC10b disturbs cristae architecture of mitochondria and reduces the life-span and fertility of flies. Moreover, using fluorescence nanoscopy and electron tomography, we demonstrate that despite its high similarity to the MIC10 proteins from yeast and humans, DmMIC10b exhibits the unique ability to polymerize into elongated filaments upon overexpression. DmMIC10b filaments form bundles which accumulate in the intermembrane space and alter the shape of mitochondrial cristae membranes. We show that the formation of the filaments relies on conserved glycine and cysteine residues and is suppressed by co-expression of other MICOS proteins. Thereby, our findings provide new insights into the regulation of MICOS in flies.

## Introduction

Mitochondria are organelles that are key for the energy metabolism of all eukaryotic cells. They rely on a double-membrane structure to generate the ubiquitous energy carrier adenosine triphosphate (ATP) by oxidative phosphorylation (OXPHOS). The two mitochondrial membranes exhibit distinct protein and lipid compositions and a strikingly different topology. The smooth outer membrane (OM) surrounds the organelle and mediates import of metabolites and proteins. The inner membrane (IM) exhibits a much larger surface and is functionally and structurally subdivided into two domains. The inner boundary membrane (IBM) parallels the OM, whereas the cristae membrane (CM) forms well-organized invaginations, termed cristae, which point toward the interior of the organelle. The cristae are connected to the IBM by small openings, referred to as crista junctions (CJs), which feature a diameter of around 20-30 nm (Mannella, Marko, & Buttle, 1997). Proper folding of the IM into cristae is closely related to the function of mitochondria and a disturbed mitochondrial ultrastructure has been associated with human diseases, including neurodegenerative disorders, metabolic diseases, and cardiomyopathies (Chan, 2012; Friedman & Nunnari, 2014; Nunnari & Suomalainen, 2012; Pernas & Scorrano, 2016; Suomalainen & Battersby, 2018; Wai & Langer, 2016).

The mitochondrial contact site and cristae organizing system (MICOS) is a large hetero-oligomeric complex situated in the IM (Harner et al., 2011; Hoppins et al., 2011; von der Malsburg et al., 2011). It is highly conserved across protozoa, fungi and animals (Wideman & Muñoz-Gómez, 2018) and consists of at least seven different proteins (MIC60, MIC27, MIC26, MIC25, MIC19, MIC13 and MIC10) in humans (Mukherjee, Ghosh, & Meinecke, 2021; van der Laan, Horvath, & Pfanner, 2016). MICOS is a crucial determinant for the biogenesis and maintenance of cristae and CJs. The holo-MICOS complex consists of two sub-complexes, namely the MIC60-subcomplex and the MIC10-subcomplex. The MIC60-subcomplex, composed of MIC60, MIC25 and MIC19, is critically important for the stability of MICOS and the formation of CJs. Loss of MIC60 causes degradation of all MICOS proteins, disrupts virtually all CJs and strongly disturbs the cristae organization in yeast and human cells (Alkhaja et al., 2012; Harner et al., 2011; Kondadi et al., 2020; Li et al., 2016; T. Stephan et al., 2020). The MIC10-subcomplex, consisting of MIC10, MIC13, MIC26 and MIC27 in humans, is crucial for a proper cristae architecture and influences the distribution of CJs. MIC10, a core subunit of this MICOS-subcomplex, is a small, highly conserved transmembrane (TM) protein of about 70-100 amino acids, depending on the species. Biochemical studies of the budding yeast ScMIC10 suggest that MIC10 features a hairpin-like structure with its N- and C-termini pointing toward the IMS. Each of the two TM domains of MIC10 contain a highly conserved glycine-rich motif (Barbot et al., 2015; Bohnert et al., 2015). For ScMIC10 it has been shown that these glycine-rich motifs mediate the formation of stable oligomers that are suggested to bend the mitochondrial IM to support cristae formation (Barbot et al., 2015; Bohnert et al., 2015).

MICOS is best studied in budding yeast and human cells, whereas it is less well characterized in the fly *Drosophila melanogaster*. Depletion of the MIC60 orthologue Mitofilin/Dmel_CG6455 results in aberrant cristae morphologies and has been related to impaired synaptic functions and leading to the death of flies in the late pupal stage (Tsai, Papakyrikos, Hsieh, & Wang, 2017). Likewise, depletion of the MIC13 orthologue QIL1/Dmel_CG760 or of the MIC26-MIC27 orthologue Dmel_CG5903 resulted in altered mitochondrial morphology and aberrant cristae (Guarani et al., 2015; Wang, Hsu, Lin, & Fu, 2020). These morphological changes were associated with reductions in climbing activity, indicating deficits in muscle function (Wang et al., 2020).

Genome sequencing (FlyBase, release FB2022_06) suggested the existence of three MIC10 orthologues in *D. melanogaster* (Dmel_CG12479/MINOS1a, Dmel_CG41128/MINOS1b and Dmel_CG13564/MINOS1c) (Gramates et al., 2022; Pfanner et al., 2014). Their functional role has so far not been reported in any detail. In this study, we show that the ubiquitously expressed Dmel_CG41128 is the major MIC10 orthologue in flies which we refer to as DmMIC10b. Loss of DmMIC10b disturbs mitochondrial ultrastructure and reduces the life span of flies. Overexpression of DmMIC10b leads to the formation of long cristae-shaping filaments along the IMS. We demonstrate that this striking behavior of DmMIC10b relies on several conserved amino acid residues and can be efficiently suppressed by co-expression of DmMIC13 or DmMIC26, but not by their human orthologues. The findings provide new insights into the regulation of MIC10 oligomerization.

## Material and Methods

### Plasmids

#### Plasmids for generation of DmMIC10b knockout

Guide RNA (gRNA) sequences upstream of the CG41128 5’UTR and downstream of the CG41128 3’UTR were identified using the web-based tool introduced by Gratz et al, 2014 (http://tools.flycrispr.molbio.wisc.edu/targetFinder/).

Criteria were as follows: CRISPR targets with 5’ G; Stringency: High; PAM: NGG Only and the closest site, with no off target to the UTR. The respective sequences were purchased as 5’-phosphorylated oligonucleotides, annealed and ligated into the BbsI sites of pU6-BbsIchiRNA (Addgene 45946) (Gratz et al. 2013a) to create ***pU6-BbsI-chiRNA-CG41128-5’gRNA*** (oligonucleotides 1/2) and ***pU6-BbsI-chiRNA-CG41128-3’gRNA*** (oligonucleotides 3/4).

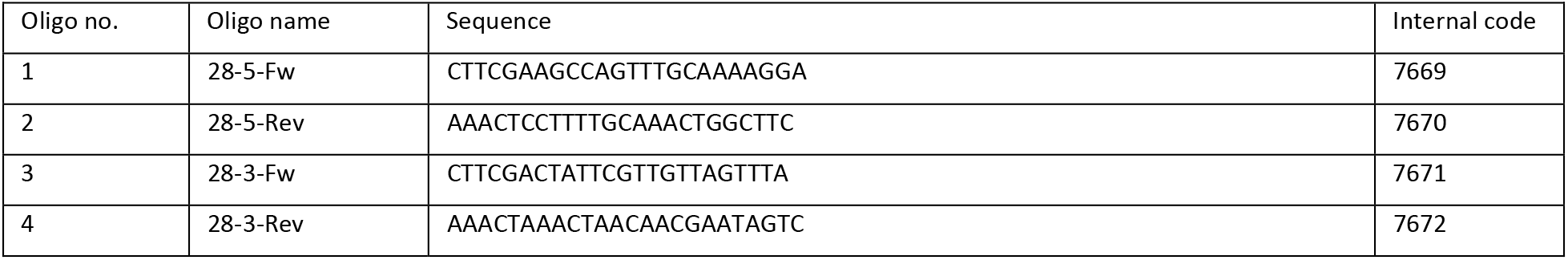

The donor plasmid ***pHD-attp-DsRed-CG41128*** for homologous recombination was generated using Gibson assembly. To this end, pHD-DsRed-attP (Addgene 51019) (Gratz et al., 2014) was double-digested with NotI/XhoI, and used as the vector backbone. Genomic DNA of a wild type strain (w^-^) was used as a template to amplify homology arms of about 1 kbp flanking the cleavage sites, and the missing base pairs from the cleavage site to the corresponding UTR beginning/end. The DsRed-attp fragment was amplified from pHD-DsRed-attp. The NEBuilder Assembly Tool was used to design the oligonucleotides and Gibson Assembly Master Mix (NEB) for the assembly reaction.

The 5’-homology arm was amplified using the oligonucleotides 5/6. The 3’-homology arm was amplified using the oligonucleotides 13/14. The 5’-missing arm was amplified using the oligonucleotides 7/8. The 3’-missing arm was amplified using the oligonucleotides 11/12. The DsRed-attp fragment was amplified using the oligonucleotides 9/10.

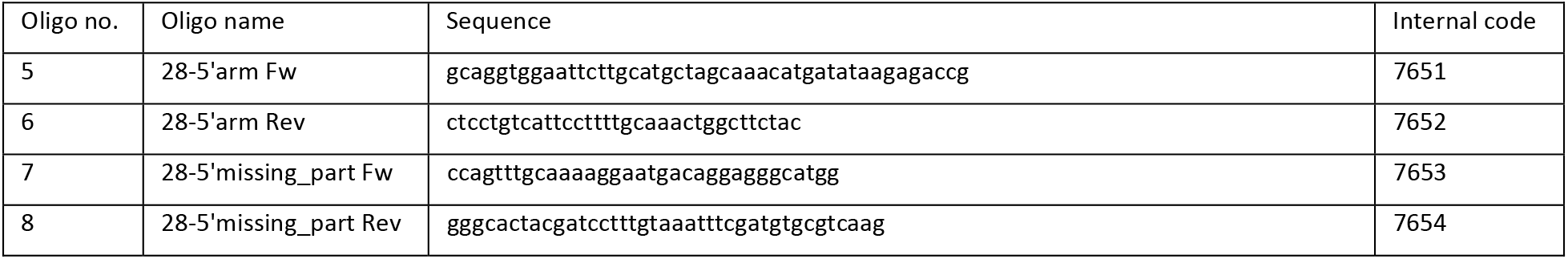
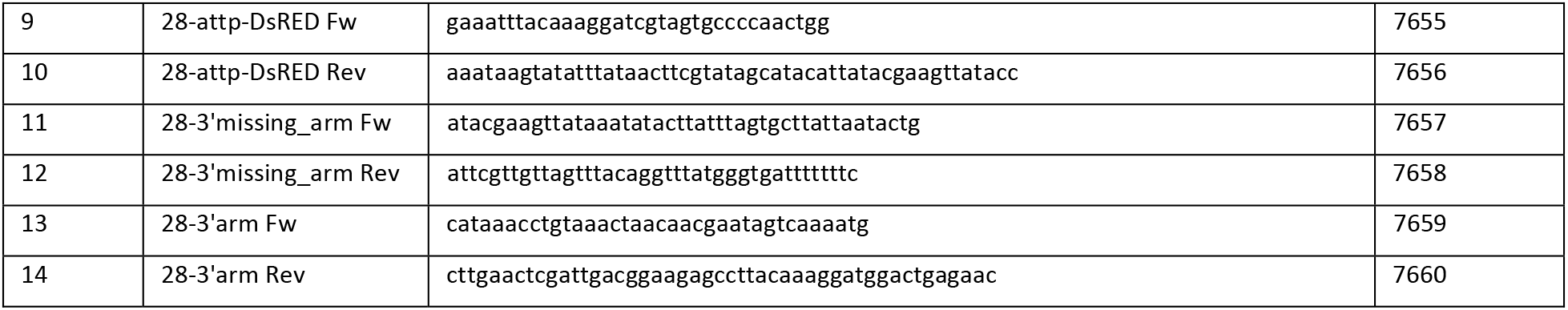

### Expression plasmids for S2 cells

#### pUAS-CG41128

DmMIC10b was amplified from cDNA using the oligonucleotides 15/16 and ligated into the EagI/XhoI restrictions sites of pUASPattB (a gift from Alf Herzig, MPI for Biophysical Chemistry, Goettingen).

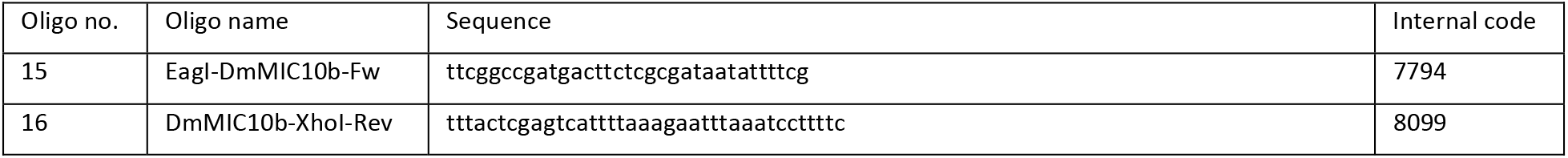

#### pUAS-CG41128-FLAG

DmMIC10b-FLAG was amplified from cDNA using the oligonucleotides 15/17 and ligated into the EagI/XhoI restrictions sites of pUASPattB.

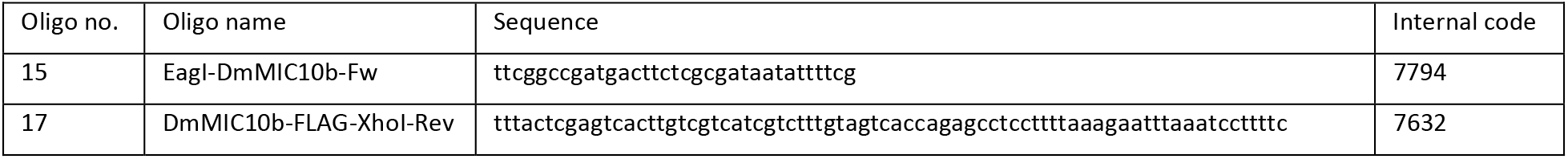

#### pUAS-CG41128(G52L/G56L)-FLAG and pUAS-CG41128(G54L/G58L)-FLAG

Glycine mutants were produced by site-directed mutagenesis of pUASPattB-CG41128-FLAG using the oligonucleotides 18/19 or 20/21, respectively.

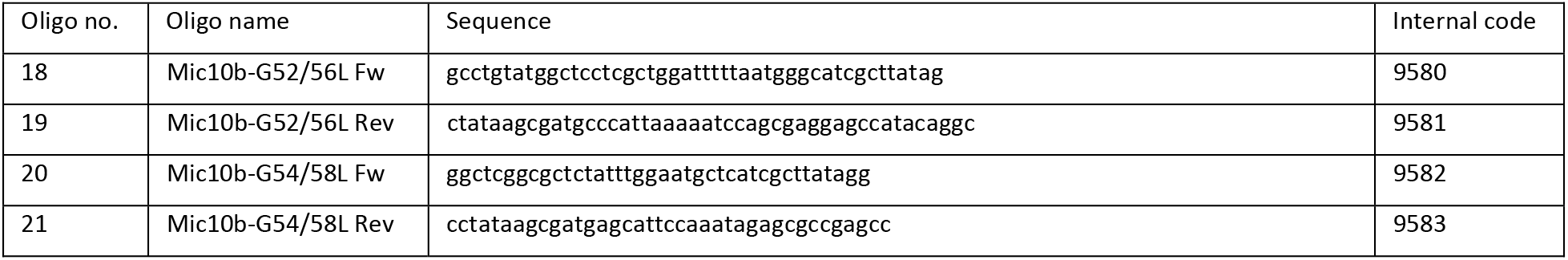

#### pUAS-HsMIC10-FLAG

HsMIC10-FLAG was amplified by PCR using the oligonucleotides 22/23 and ligated into the EagI and XhoI restriction sites of pUASPattB.

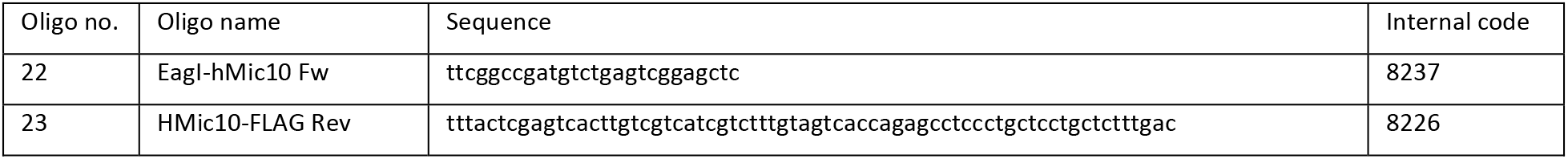

#### pUAS-ScMIC10-FLAG

ScMIC10-FLAG was amplified by PCR using the oligonucleotides 24/25 and integrated into the EagI and XhoI restriction sites of pUASPattB.

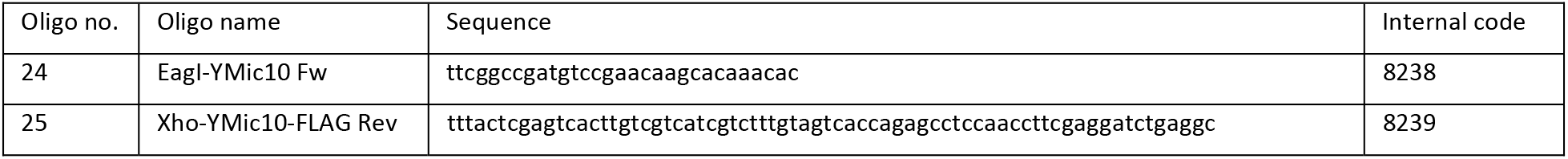

### Expression plasmids for mammalian cells

#### pcDNA3.1-DmMIC10b-FLAG

Humanized DmMIC10b-FLAG (GenScript, Piscataway Township, NJ, USA) was amplified by PCR using the nucleotides 26/27 and inserted into the HindIII and XhoI restriction sites of pcDNA3.1(+) (Thermo Fisher Scientific). The point mutants MIC10b(C13S)-FLAG, DmMIC10b(C19S)-FLAG, DmMIC10b(C28S)-FLAG, DmMIC10b(I41F)-FLAG, DmMIC10b(C64S)-FLAG were produced using the Site-Directed Mutagenesis Kit (NEB) and the oligonucleotides given below.

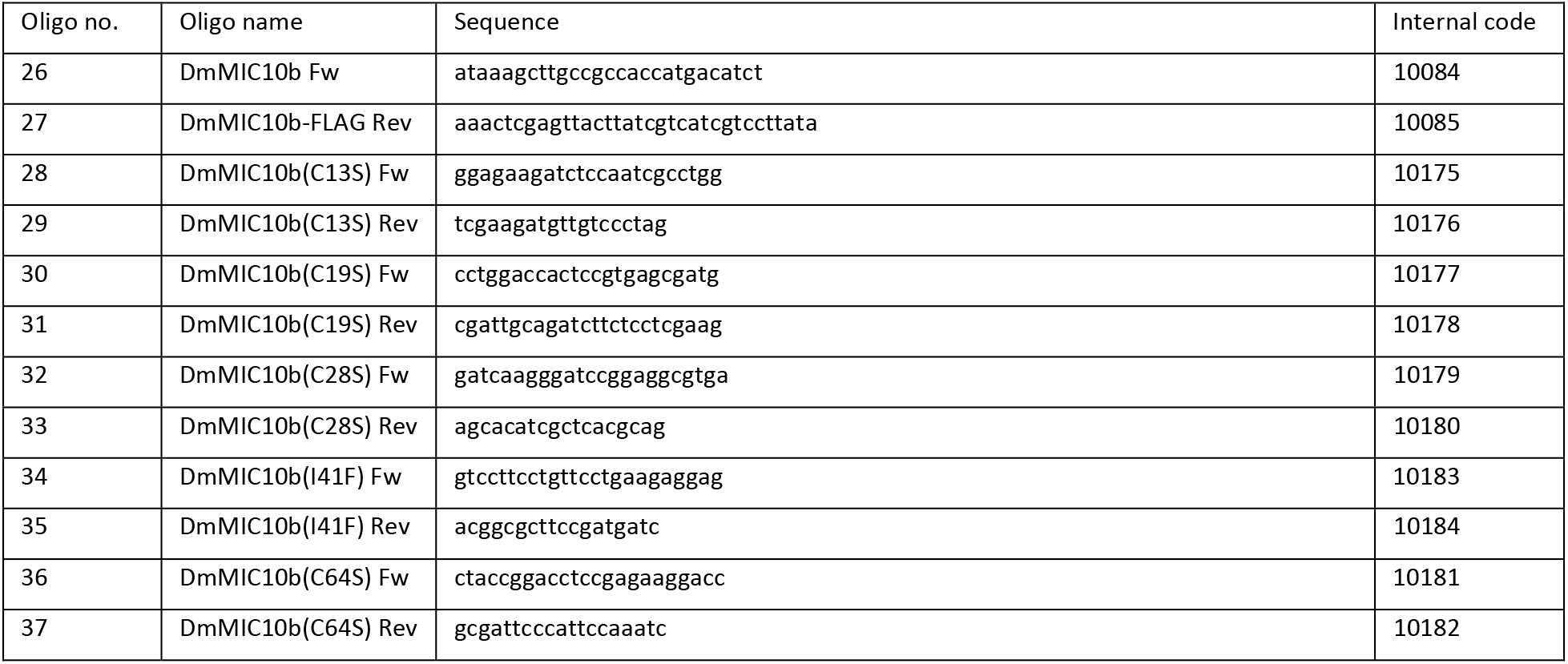

#### AAVS1-TRE3G-DmMIC10-FLAG-T2A-EGFP

Humanized DmMIC10b-FLAG flanked by AgeI and EcoRV restriction sites (obtained from GenScript) was inserted into the AgeI/EcoRV restriction sites of AAVS1-TRE3G-Mic10-FLAG-T2A-EGFP (T. Stephan et al., 2020).

#### AAVS1-TRE3G-DmMIC10-T2A-EGFP

TRE3G-DmMIC10 was amplified (including SpeI and AgeI restriction sites) from AAVS1-TRE3G-DmMIC10b-FLAG-T2A-EGFP by PCR using the oligonucleotides 38/39. AAVS1-TRE3G-Mic10-FLAG-T2A-EGFP (T.

Stephan et al., 2020) was linearized using SpeI and AgeI restriction endonucleases and the PCR product was ligated into the plasmid to remove the FLAG-epitope.

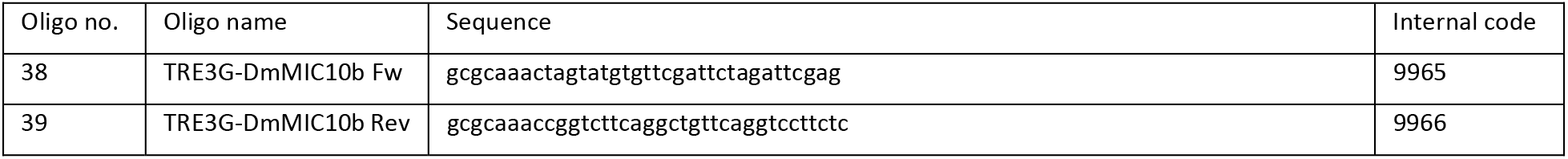

#### pFLAG-CMV5.1-ScMIC10-FLAG

ScMIC10-FLAG was amplified by PCR using the oligonucleotides 40/41 and integrated into the EagI/XbaI restriction sites of pFLAG-CMV5.1 (Sigma Aldrich).

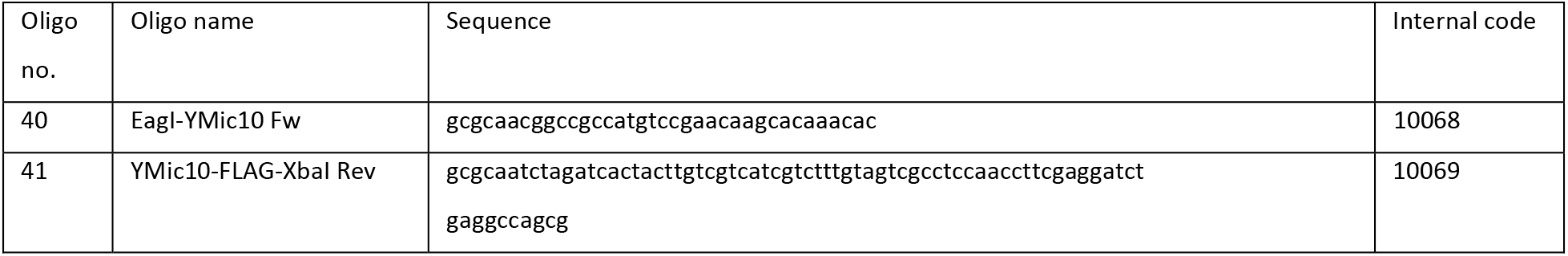

### Co-Expression of DmMIC10b-FLAG and Spot-tagged MICOS proteins

#### DmMic13-(C90S)-Spot_pJET1.2

The substitution mutation DmMIC13(C90S)-Spot was introduced to DmMIC13-Spot by site-directed PCR mutagenesis. To this end, DmMIC13-Spot (GenScript) was amplified by PCR using the oligonucleotides 42/43 to introduce the substitution mutation C90S to DmMIC13-Spot by site-directed PCR mutagenesis.

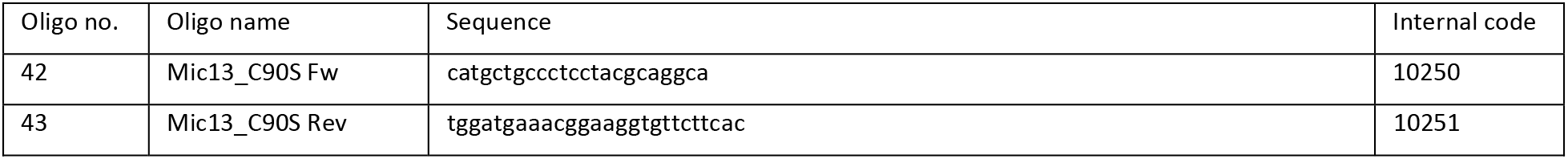

#### AAVS1-TRE3G-DmMIC10b-FLAG-T2A-DmMIC13-Spot and AAVS1-TRE3G-DmMIC10b-FLAG-T2A-DmMIC13(C90S)-Spot

DmMIC13-Spot or DmMIC13(C90S)-Spot were amplified by PCR using the oligonucleotides 44/45. AAVS1-TRE3G-DmMIC10b-FLAG-T2A-EGFP was linearized using SalI/MluI restriction endonucleases and DmMIC13-Spot or DmMIC13(C90S)-Spot were ligated into the backbone, respectively.

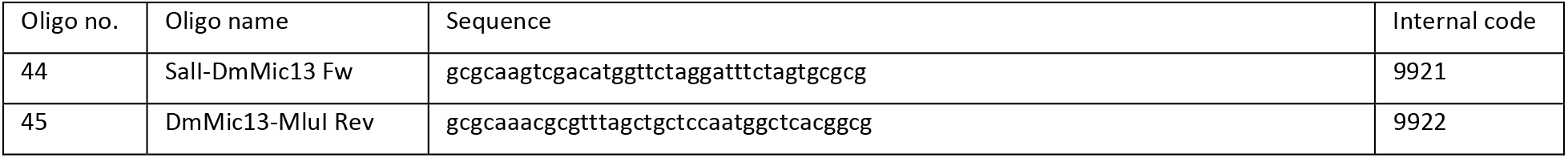

#### AAVS1-TRE3G-DmMIC10b-FLAG-T2A-HsMic13-Spot

HsMIC13 was amplified by PCR using the oligonucleotides 46/47. AAVS1-TRE3G-DmMIC10b-FLAG-T2A-EGFP was linearized using SalI/MluI restriction endonucleases and HsMIC13-Spot was ligated into the backbone.

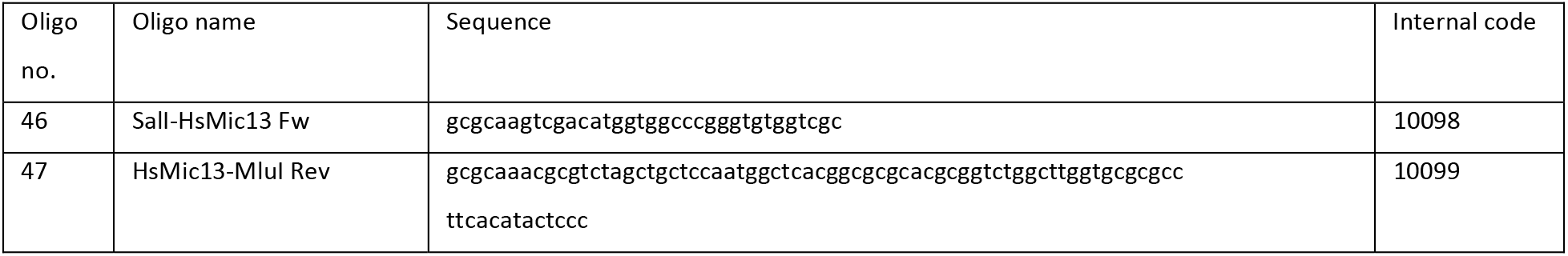

#### AAVS1-TRE3G-DmMIC10b-FLAG-T2A-DmMIC26-Spot

DmMIC26-Spot (GenScript) was amplified by PCR using the oligonucleotides 48/49. AAVS1-TRE3G-DmMIC10b-FLAG-T2A-EGFP was linearized using SalI/MluI restriction endonucleases and DmMIC26-Spot was ligated into the backbone.

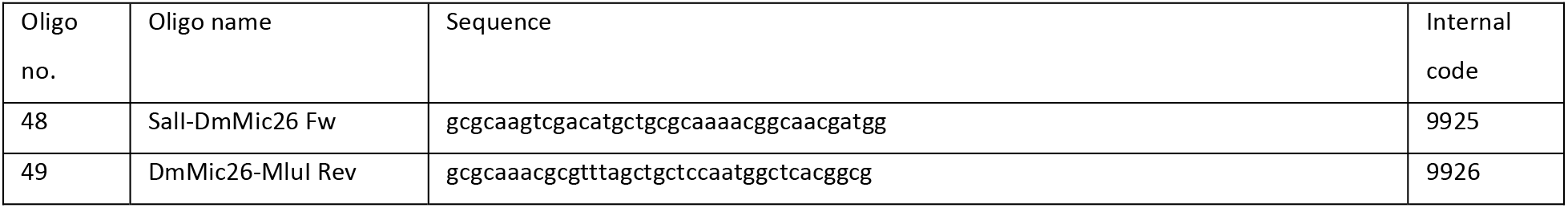

#### AAVS1-TRE3G-DmMIC10b-FLAG-T2A-DmMIC19-spot

DmMIC19-Spot (GenScript) was amplified by PCR using the oligonucleotides 50/51. AAVS1-TRE3G-DmMIC10b-FLAG-T2A-EGFP was linearized using SalI/MluI restriction endonucleases and DmMIC19-Spot was ligated into the backbone.

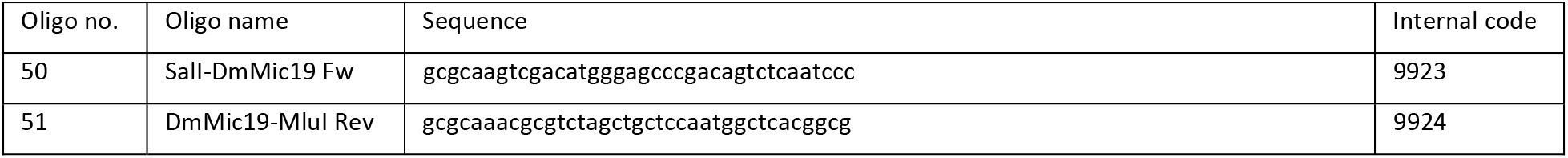

### Expression plasmids for bacteria

#### pPROEXHTb-DmMIC10b and derived point mutants

DmMIC10b cDNA was amplified by PCR using oligonucleotides 52/53 and integrated into pPROEXHTb (Thermo Fisher) via the NcoI/EcoRI restriction sites. Point mutants were generated by site-directed genesis PCR.

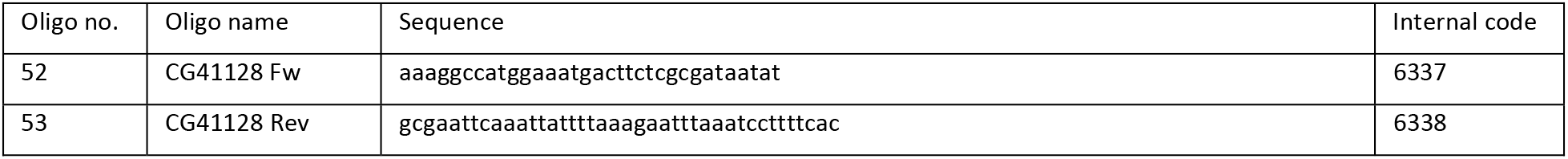

### Purification of DmMIC10b from *E. coli* and preparation for EM

The purification of His-tagged DmMIC10b was performed as described previously (Barbot et al., 2015). In brief, *E. coli* BL21 (DE3) cells were collected by centrifugation after expression of His_6_-DmMIC10b (1 mM isopropyl-β-D-thiogalactopyranoside (IPTG), 3h, 37°C) and stored at -20°C until purification. After thawing, cells were lysed, inclusion bodies were isolated and subsequently dissolved in resuspension buffer containing 8 M urea, 150 mM NaCl, 20 mM Tris-HCl, 40 mM imidazole, 0.5 mM DTT, pH 8.0. The mixture was applied to a HisTrap column (5 ml) and eluted with resuspension buffer supplemented with 500 mM imidazole. Isolated Mic10 was further subjected on HiLoad 16/600 Superdex 200 size exclusion column (GE Healthcare, Piscataway, NJ, USA). Separated fractions were analyzed by SDS-PAGE and Coomassie brilliant blue staining.

### Expression of DmMIC10b mutants in E. coli and analysis by western blotting

DmMIC10b mutants were expressed in *E. coli* BL21 (DE3). Cells were collected by centrifugation after expression (1 mM isopropyl-β-D-thiogalactopyranoside (IPTG), 3h, 37°C) and washed in salt buffer containing 150 ml NaCl, 10 mM HEPES, pH 7.4. The pellet was resuspended in lysis buffer containing 50 mM Tris (pH 8.0), 150 mM NaCl, 0.1 mg/ml lysozyme, 1 mM MgCl_2_ and cOmplete protease inhibitor (Merck), and the cells were homogenized by sonication. The homogenate was supplemented with benzonase (Sigma Aldrich) and stirred for 30 min at 4 °C. Following centrifugation, the pellet was washed with Triton X-100 wash buffer containing 50 mM Tris, 150 mM NaCl, 1 mM EDTA, 2% (w/v) Triton X-100 (pH 8.0) followed by washing with a wash buffer containing 50 mM Tris, 100 mM NaCl, 10 mM DTT, 1 mM EDTA (pH 8.0). The pellet was dissolved with 8% (w/v) sarcosyl and 1 M urea (in 50 mM Tris, pH 8.0) and insoluble material was removed by centrifugation. The solution was diluted with solubilization buffer (20 mM Tris, 150 mM NaCl, pH 8.0) to a final concentration of 1.5 % (w/v) sarcosyl, supplemented with 0.1 % (w/v) DDM and dialyzed against solubilization buffer overnight. Samples were analyzed by SDS-PAGE followed by western blotting. His-tagged DmMIC10b was detected using an anti-His antibody (Qiagen N.V., Venlo, The Netherlands, 34660).

### Cultivation of flies and life-span assay

Flies were maintained on standard fly food with cornmeal, yeast, and agar at 25°C on a 12-hour/12-hour light/dark cycle. Fly stocks used were *DmMIC10b[KO]* (this study), *w[1118]* (Bloomington Drosophila Stock center, BDSC #6326), and OR-R (BDSC #5). Wild-type control flies used were the heterozygous progeny of *w[1118]* males and OR-R females.

Life-span assay was performed using 4 biological replicates per genotype. 15 male and 15 female young flies were placed in each vial. Every 2-4 days, flies were transferred to new food vials, and both living and dead flies were counted for consistency. Counting was continued until all flies had died.

### Generation of knockout flies

*Dmel_CG41128[KO]* flies were generated by a commercial transformation service (BestGene Inc, Chino Hills, CA, USA). The injection stock was RRID:BDSC_55821 (Bloomington Drosophila Stock Center, Bloomington, IN, USA).

### Cell culture

*Drosophila* S2 cells (Thermo Fisher Scientific, Waltham, MA, USA, Gibco®, Cat. No. R69007, Lot No. 2082623) were cultivated in Schneider’s *Drosophila* Medium (Merck, Darmstadt, Germany /Sigma Aldrich, St. Louis, MO, USA, Gibco™, Cat. No. 21720024) supplemented with 1mM sodium pyruvate (Merck, Darmstadt, Germany /Sigma Aldrich, St. Louis, MO, USA, Cat. No. S8636) and 10% (v/v) fetal bovine serum (Bio & Sell GmbH, Feucht bei Nürnberg, Germany, FBS superior stable, Cat. No. FBS. S 0615). Cells were cultivated at 28 °C and ambient CO_2_ levels. Kidney fibroblast-like cells (COS-7) from the African green monkey *Cercopithecus aethiops* (Merck, Darmstadt, Germany/Sigma Aldrich, St. Louis, MO, USA, Cat. No. 87021302, Lot No. 05G008) as well as human cervical cancer cells (HeLa) were cultivated in DMEM, containing 4.5 g/L Glucose and GlutaMAX^TM^ additive (Thermo Fisher Scientific, Waltham, MA, USA, Cat. No. 10566016) supplemented with 1mM sodium pyruvate (Merck, Darmstadt, Germany /Sigma Aldrich, St. Louis, MO, USA, Cat. No. S8636) and 10% (v/v) fetal bovine serum at 37 °C and 5 % CO_2_. Human osteosarcoma cells (U-2 OS) (ECACC, Porton Down, Salisbury, UK; Cat no. 92022711, Lot. 17E015) were cultivated at 37 °C and 5 % CO_2_ and grown in McCoýs medium (Thermo Fisher Scientific, Waltham, MA, USA) supplemented with 10 % (v/v) fetal bovine serum, and 1 % (v/v) sodium pyruvate.

### Transfection of HeLa, COS-7 and S2 cells

S2 cells were split the day before transfection. On the day of transfection 1.5 x 10^6^ cells were seeded per well of a 6-well dish. Transient transfection was achieved using Effectene transfection reagent (Qiagen, Hilden, Germany, Cat. No. 301425) and 2 µg of plasmid DNA per well. After 23 h the cells were detached and reseeded on glass cover slips coated with Concanavalin A (Merck, Darmstadt, Germany, /Sigma Aldrich, St. Louis, MO, USA, Cat. No. C5275) for one hour.

Mammalian cells were seeded on cover slips (for light microscopy), aclar film (for electron microscopy) and in cell culture dishes (for biochemistry) one day prior to transfection. Transient transfection was achieved by jetPRIME^®^ transfection reagent (Polyplus-transfection, Illkirch-Graffenstaden, France, Cat. No. 114-15) following the manufacturer protocol.

### Generation of antibodies against DmMICOS proteins

*Drosophila* anti-MICOS polyclonal antibodies were produced by injecting purified protein or synthetic peptide into rabbits (Prof. Hermann Ammer, München, Germany). For anti-MIC10 antibody generation, recombinantly expressed and affinity-purified MIC10 protein and for anti-MIC60 and anti-MIC13 antibody generation AAKPKDNPLPRDVVELEKA and GDSDQTDKLYNDIKSELRPH synthetic peptides were injected into rabbits, respectively. All antibodies were affinity purified.

### Mitochondrial isolation

Mitochondria were isolated as described previously (Panov & Orynbayeva, 2013) with slight modifications. Flies were taken up in cold TH-buffer, containing 300 mM Trehalose, 10 mM KCl, 10 mM HEPES (pH 7.4), 2 mM PMSF, 0.1 mg BSA/ml and protease inhibitor mix (Roche, 04693116001). The flies were then two times homogenized with a dounce homogenizer (800 rpm/min) and large remaining fragments were pelleted at 400 x g for 10 min at 4°C. Afterwards, remaining pieces were removed by centrifugation (800 x g, 8 min, 4°C) and the mitochondria containing supernatant saved in new vial. To collect mitochondria a further spin at 11,000 x g, 10 min, 4°C was performed, pellets from separate reaction tubes pooled and washed with BSA-free TH-buffer. The mitochondria concentration was determined using Bradford assay.

### Affinity purification from fly mitochondria

Isolated mitochondria (1 mg) were transferred to lysis buffer (150 mM NaCl, 10% glycerol (v/v), 20 mM MgCl_2_, 2 mM PMSF, 50 mM Tris-HCl, pH 7.4, 1% digitonin (v/w), protease inhibitor (Roche 04693116001)) and agitated 30 min at 4 °C. Debris were removed by centrifugation (15 min, 16,000 x g, 4°C) and the cleared supernatant was transferred Proteine-A sepharose conjugated with anti-DmMIC60 or control antisera. After 1 h binding at 4°C, beads were washed 10 times with washing buffer (50 mM Tris-HCl, pH 7.4, 150 mM NaCl, 10% glycerol (v/v), 20 mM MgCl_2_, 1 mM PMSF, 0.3% digitonin (v/w)). Proteins were eluted by addition of 0.1 M glycine pH 2.8 at room temperature (RT) for 5 min. Eluates were neutralized and analyzed by SDS-PAGE and immunoblotting using specific antibodies against DmMIC60 (this study), DmMIC10b (this study), DmMIC13 (this study), NDUFA9, and RIESKE (Dennerlein et al., 2021).

### Affinity purification from cell lysates

COS-7 and HeLa cells were seeded in 15 cm cell culture dishes and cultured overnight. Cells were transfected with AAVS1-TRE3G-Mic10-FLAG-T2A-EGFP (Stephan et al., 2020, EMBOJ), AAVS1-TRE3G-DmMIC10-FLAG-T2A-EGFP, or AAVS1-TRE3G-EGFP (Qian et al., 2014) (Addgene 52343), respectively. Medium was exchanged 4h after transfection and expression was induced by adding 1 µg/ml doxycycline hyclate for 24 h. Cells were harvested by trypsinization and centrifugation. All following steps were performed at 4 °C. The pellet was washed with PBS and resuspended in 1.5 ml lysis buffer containing 20 mM Tris (pH 7.0), 100 mM NaCl, 1 mM EDTA, 10 % (v/v) glycerol, 1% (w/v) digitonin (Carl Roth GmbH, Karlsruhe, Germany) and cOmplete protease inhibitor (Merck). Samples were rotated on a wheel for 1h. After centrifugation at 13,000xg for 10 min, the supernatant was transferred onto equilibrated magnetic anti-FLAG M2 magnetic beads (Merck). Samples were rotated on a wheel for 1.75 h. The supernatant was removed and beads washed 10 times with wash buffer containing 20 mM Tris, 100 mM NaCl, 1 mM EDTA, 5% (v/v) glycerol, 0.25% (w/v) digitonin and complete protease inhibitor (pH 7.0). Elution was performed by adding 0.1 M glycine (pH 2.8) and shaking the samples at 1200 rpm for 20 min. Eluates were neutralized and analyzed by SDS-PAGE and immunoblotting using specific antibodies against FLAG (Sigma Aldrich, F3165), MIC13 (Sigma Aldrich, SAB1102836), MIC19 (Atlas Antibodies, HPA042935), COX3 (Proteintech, 55082-1-AP), MIC60 (Proteintech, 10179-1-AP) and MFN1 (Abcam, ab57602).

### Preparation of fly tissues for fluorescence microscopy

Adult testes were dissected in ice-cold PBS, then fixed at RT using 4% formaldehyde in PBT (PBS + 0.2% (v/v) Triton-X 100) for 20 minutes. After fixation, testes were rinsed in PBT (multiple rinses for a minimum of 30 total minutes), then blocked using PBTB (PBT + 0.2% (w/v) bovine serum albumin + 5% (v/v) normal goat serum) for 30 minutes. After blocking, samples were incubated in primary antibody diluted in PBTB overnight at 4°C. Then, primary antibody was removed, samples were rinsed in PBT for 30 minutes, then for 30 minutes in PBTB. Secondary antibodies diluted 1:500 in PBTB were added to the samples overnight at 4°C. Samples were then rinsed again with PBT for 30 minutes, treated with DAPI for 10 minutes to stain DNA, and finally rinsed again with PBT. Samples were stored in Vectashield (Vector Laboratories, Newark, CA, USA) until being mounted on slides and imaged. Primary antibodies used for staining testes were rabbit anti-DmMIC10b (1:3000, this work) and mouse anti-ATP5A (1:300, Abcam, ab14748).

### Sample preparation of fixed S2 cells for STED nanoscopy

Fixation and labeling of S2 cells were done as described previously (Wurm, Neumann, Schmidt, Egner, & Jakobs, 2010). In brief, cells were fixed using an 8 % (w/v) formaldehyde solution, permeabilized by incubation with a 0.25 % (v/v) Triton X-100 solution and blocked with a 5 % (w/v) bovine serum albumin (BSA) solution. Proteins of interest were labelled with antisera against the FLAG tag (Thermo Fisher Scientific, Waltham, MA, USA) and ATP5A (Abcam, Cambridge, UK) respectively. Detection was achieved via secondary antibodies custom-labelled with Alexa Fluor 594 or STAR RED, respectively.

### Sample preparation of fixed HeLa and COS-7 cells for STED nanoscopy and 4Pi STORM

Cells were transfected with pcDNA3.1-DmMIC10b-FLAG or with AAVS1-TRE3G-DmMIC10b-FLAG-T2A-EGFP. In case of the latter construct, cells were induced with 1 µg/ml doxycycline. Fixation and labeling were performed 24 h after transfection or induction, respectively (Wurm et al., 2010). In brief, cells were fixed using 4 % (w/v) or 8 % (w/v) formaldehyde solution, permeabilized by incubation with a 0.25 % (v/v) Triton X-100 solution and blocked with a 5 % (w/v) bovine serum albumin (BSA) solution. Proteins of interest were labelled with antibodies against DmMIC10b (this study), FLAG tag (Merck, Darmstadt, Germany /Sigma Aldrich, St. Louis, MO, USA), Spot tag (ChromoTek, Planegg-Martinsried, Germany), TOMM22 (Merck, Darmstadt, Germany /Sigma Aldrich, St. Louis, MO, USA) and MIC60 (Abcam) respectively. For STED nanoscopy, detection was achieved via secondary antibodies custom-labelled with Alexa Fluor 594 or STAR RED, respectively. Cells were mounted using Mowiol containing 1,4-Diazabicyclo[2.2.2]octan (DABCO).

For 4Pi-STORM, samples were prepared on 18-mm diameter glass coverslips which were coated over one quarter of their surface with a reflective aluminum layer, which was used for initial alignment of the sample within the 4Pi-STORM microscope. Cells were plated on the coated side of the coverslip, and fluorescently labeled as described before (Bates et al., 2022). For two-color imaging, cells were fixed and stained with antibodies against MIC60 (Abcam) and the FLAG epitope (Sigma Aldrich). Primary antibodies were detected with Fab fragments coupled to Alexa Fluor 647 (Thermo Fisher) or antibodies labeled with Cy5.5 (Jackson ImmunoResearch, Cambridge, UK). Prior to imaging, the sample was washed with PBS and mounted in STORM imaging buffer in a sandwich configuration with a second glass coverslip. The two coverslips were sealed together around their edge with fast-curing epoxy glue (Uhu GmbH, Bühl, Germany).

### Sample preparation for live-cell STED nanoscopy

HeLa and COS-7 cells were seeded in glass bottom dishes (ibidi GmbH, Gräfelfing, Germany) and grown at 37°C and 5% CO_2_ overnight. Cells were co-transfected with AAVS1-TRE3G-DmMIC10b-FLAG-T2A-EGFP and AAVS1-Blasticidin-CAG-COX8A-SNAP (Till Stephan, Roesch, Riedel, & Jakobs, 2019). 4 hours after transfection, the culture medium was exchanged and the expression of DmMIC10b-FLAG-T2A-EGFP was induced by adding 1 µg/ml doxycycline hyclate (Sigma Aldrich, St. Louis, MO, USA) to the growth medium for 24h. Prior to STED imaging, cells were stained with 1 µM SNAP-cell SiR-647 (NEB) for 30 min at 37 °C and 5 % CO_2_. Transfected cells were identified based on the cytosolic EGFP reporter and recorded by live-cell STED nanoscopy at RT.

### Sample preparation of cultivated cells for electron tomography

Aclar®33C disks were punched with 18 mm diameter using 0.198 mm thick aclar film (Plano, Wetzlar, Germany) and sterilized with 70% ethanol before usage. On these, COS-7 cells or MICOS-depleted HeLa cells (T. Stephan et al., 2020) were grown were grown to ∼ 70% confluency and transfected with pcDNA3.1-DmMIC10b-FLAG. 24h after transfection, samples were immobilized and fixed with 2.5% (v/v) glutaraldehyde in 0.1 M cacodylate buffer at pH 7.4, first for 1 h at RT, then transferred to 4 °C until the next day. Before post-fixation, the cells were additionally stained with 1% osmium tetroxide and 1.5% (w/v) K_4_[Fe(CN)_6_] in 0.1 M cacodylate buffer at pH 7.4 for 1 h at RT. Following post-fixation in 1% osmium tetroxide for 1 h at RT and pre-embedding en-bloc staining with 1% (w/v) uranyl acetate for 30 min at RT, samples were dehydrated and embedded in Agar 100 resin (Plano, Wetzlar, Germany).

### Sample preparation of fly tissues for electron microscopy

After extraction, fly brains were fixed in bulk by immersion with 2% glutaraldehyde in 0.1 M cacodylate buffer at pH 7.4. Fixation was completed overnight at 4 °C. After three washing steps with 0.1 M cacodylate buffer the brains were stained with 1% (w/v) osmium tetroxide for 1 h at RT followed by en-bloc staining with 1% (w/v) uranyl acetate for 30 min at RT. Subsequently the brains were dehydrated in a graded series of ethanol and finally flat-embedded in a thin layer of Agar100 resin in between two Aclar sheets. Individual brains were manually cut out of the resin with a saw and 80 nm thick sections were collected on Formvar coated copper grids. 10x10 tiles at 8600x original magnification were recorded at randomly selected areas of the fly brains.

### Preparation of recombinant DmMIC10b for negative stain EM

DmMIC10b purified in presence of urea was dialyzed (against 20 mM Tris, 100 mM NaCl, 2 mM DTT, pH 8.0) to precipitate DmMIC10b. The precipitate was solubilized by adding 10% (w/v) sarcosyl and 5 % (w/v) digitonin followed by thorough pipetting. The obtained solution was afterward diluted with a suspension buffer containing 20 mM Tris, 100 mM NaCl, 3% (w/v) DDM, 2mM DTT, pH 8.0 up to final concentration of 0.2% sarcosyl and 0.05% digitonin. The solution was supplemented with 10 mM imidazole and loaded onto magnetic His-Beads. Samples were rotated on a wheel for 1.5 h at 4 °C. The supernatant was removed and the beads were washed five times with a wash buffer containing 20 mM Tris, 100 mM NaCl, 10 mM imidazole, 1 mM DTT, 0.05 % (w/v) DDM, pH 8.0. Afterward, the beads were washed 5 times with a wash buffer containing 20 mM Tris, 100 mM NaCl, 1 mM imidazole, 1 mM DTT, 0.01 % (w/v) DDM, pH 8.0. Elution was performed with 500 mM imidazole. Samples were dialyzed (against 20 mM Tris, 100 mM NaCl, 1 mM DTT, 0.01% (w/v) DDM, pH 8.0) using a 5 kDa cutoff membrane. The samples were analyzed by SDS-PAGE and Coomassie Brilliant Blue staining. For negative staining EM, the obtained protein samples were bound to a glow discharged carbon foil covered 400 mesh copper grid. After successive washing with water, samples were stained with 1% (w/v) uranyl acetate aq. and evaluated at RT using a Talos L120C (Thermo Fisher Scientific).

### Fluorescence microscopy

Imaging of testes was performed on a Zeiss LSM 700 confocal laser-scanning microscope (Carl Zeiss AG, Oberkochen, Germany). Confocal light microscopy images of cultured cells were acquired using a Leica TCS SP8 confocal laser-scanning microscope (Leica Microsystems, Wetzlar, Germany) equipped with a HC PL APO 63x/1.40 Oil objective (Leica Microsystems, Wetzlar, Germany).

STED nanoscopy was performed using an Expert Line quad scanning STED microscope (Abberior Instruments, Göttingen, Germany) equipped with a UPlanSApo 100x/1,40 Oil objective (Olympus, Tokyo, Japan). Alexa Fluor 594 was excited at 561 nm wavelength and STAR RED was excited at 640 nm wavelength. STED was performed at 775 nm wavelength. SiR-647 was excited at 640 nm wavelength and STED was performed at 775 nm wavelength as described previously (Stephan et al., 2019, SciRep).

For 4Pi-STORM, cells were imaged on the 4Pi-STORM microscope by illuminating with 642nm laser light and recording the on-off blinking events of individual fluorescent molecules on a CCD camera, for approximately 100000 camera frames at a rate of 100Hz. To achieve a larger imaging depth, the sample was periodically shifted along the optical axis (z-coordinate) in steps of 500nm during the recording. Afterwards, in a post-processing step, each blinking event was analyzed to determine a 3D molecular coordinate, and the full set of coordinates plotted together in 3D space forms the 3D 4Pi-STORM image. Multicolor imaging was achieved by distinguishing the Alexa Fluor 647 and Cy5.5 according to the ratio of photons for each fluorophore detected in the four image channels of the microscope. The spatial resolution of the processed 4Pi-STORM image is approximately 10nm in all dimensions. Full details of the microscope, imagine procedure, and data analysis are described in (Bates et al., 2022).

### Electron microscopy

For 2D analysis, images of ultrathin sections of ∼70 nm thickness were recorded on a Philips CM120 BioTwin transmission electron microscope (Philips Inc., Eindhoven, the Netherlands). Usually, 2D images of at least 20 different cells were randomly recorded for each sample at 8600× original magnification using a TemCam 224A slow scan CCD camera (TVIPS, Gauting, Germany).

For electron tomography, tilt-series of thin sections of ∼ 230 nm that were additionally decorated with 10 nm gold beads on both sides were recorded on a Talos L120C transmission electron microscope (Thermo Fisher Scientific/FEI company, Hillsboro, Oregon, USA) at 17,500× original magnification using a Ceta 4k × 4k CMOS camera in unbinning mode. Series were recorded from -65.0° to 65.0° with a 3° dose-symmetric angular increase. The series were calculated using Etomo (David Mastronade, http://bio3d.colorado.edu/).

### Imaging data processing

TEM recordings of thin sections and ET data were processed in Fiji using the median filter. If not stated otherwise, STED nanoscopy data were smoothed with a lowpass filter using the Imspector software (Abberior Instruments). For deconvolution, we used the Richardson Lucy algorithm in the Imspector software. For volume renderings of 4Pi-STORM data, we relied on Imaris (Bitplane, Belfast, UK). Confocal images of fly tissue were processed using ImageJ and Adobe Illustrator. The seminal vesicle areas were calculated using the “Measure” feature in ImageJ.

## Results

### The mitochondrial protein Dmel_CG41128 is homologous to MIC10 from yeast and humans

The genome of *D. melanogaster* encodes three different proteins with noticeable sequence similarity to the MIC10 proteins from yeast (ScMIC10) and humans (HsMIC10): MINOS1a/ Dmel_CG12479; MINOS1b/ Dmel_CG41128; and MINOS1c/ Dmel_CG13564. Like MIC10 from yeast and humans, these three proteins contain two putative transmembrane domains (TMD) with conserved glycine-rich motifs (Figure 1A). Each N-terminal TM segment contains a GxxxG motif (Engelman motif), which has been reported to mediate oligomerization of several membrane proteins (Russ & Engelman, 2000).

**Figure 1:**
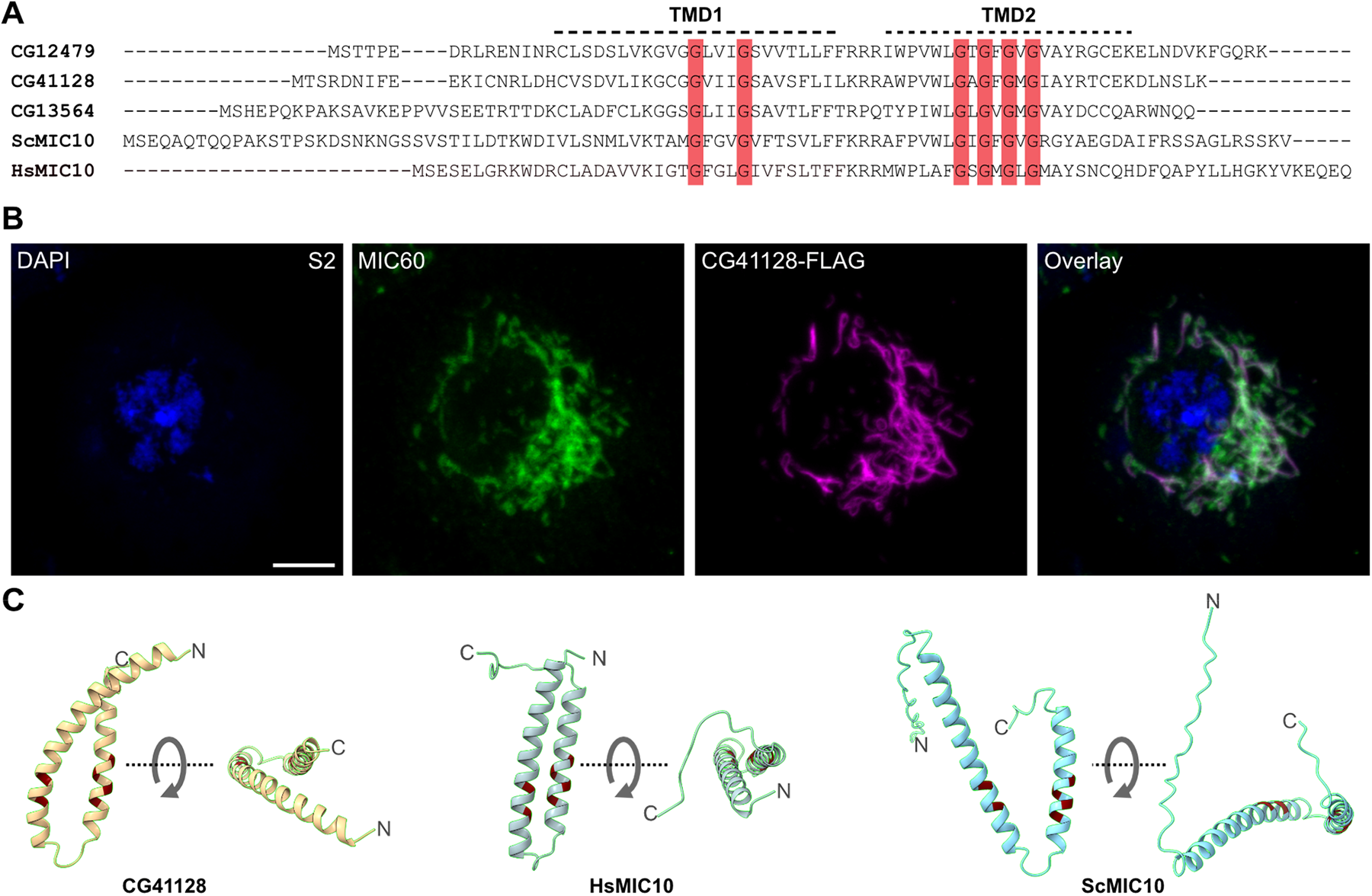
*Drosophila melanogaster* has three MIC10-like proteins. A. Sequence alignment of three putative MIC10 proteins from *D. melanogaster*, MIC10 from *S. cerevisiae* (ScMIC10) and MIC10 from H. sapiens (HsMIC10). The two putative transmembrane domains (TMD, indicated by dashed lines) and highly conserved GxxxG and GxGxGxG motifs (red shading) are highlighted. B. Immunofluorescence recording of S2 cells expressing CG41128-FLAG. C. AlphaFold2 structure predictions of Dmel_CG41128, HsMIC10 and ScMIC10. Positions of conserved glycine residues are highlighted in red. Scale bar: 5 µm.

Each C-terminal TM domain contains a related, highly conserved GxGxGxG motif, which has been shown to be crucial for the oligomerization of ScMIC10 (Barbot et al., 2015; Bohnert et al., 2015). However, only Dmel_CG41128 is ubiquitously expressed throughout all analyzed developmental stages, whereas Dmel_CG12479 and Dmel_CG13564 are testes-specific proteins (FlyBase **FB2022_06)**. Therefore, we decided to focus on the investigation of Dmel_CG41128.

Expression of Dmel_CG41128-FLAG in S2 cells and subsequent immunolabeling and fluorescence microscopy demonstrated that Dmel_CG41128 indeed localized to mitochondria (Figure 1B). We next used AlphaFold2 to predict the overall structure of Dmel_CG41128, ScMIC10 and HsMIC10. The AlphaFold2 algorithm predicted a hairpin topology reminiscent of some ER-resident reticulons (Yang & Strittmatter, 2007) for all three proteins, with the conserved GxxxG and GxGxGxG motifs being oriented to each other in a similar way (Figure 1C). The predicted structures are fully in line with previous experimental studies on ScMIC10, which demonstrated that two TM domains of ScMIC10 are linked by a short loop that points towards the matrix while the termini of the protein point towards the inter membrane space (IMS) (Barbot et al., 2015; Bohnert et al., 2015).

We conclude that Dmel_CG41128 is a mitochondrial protein that shows sequence homology with known MIC10 proteins and presumably features a comparable hairpin-like shape, making it a promising candidate for the MIC10 subunit from *D. melanogaster*.

### Loss of Dmel_CG41128 reduces the life span and the fertility of flies

In humans, the loss of subunits of the MIC10 subcomplex is associated with severe diseases such as mitochondrial encephalopathy, myopathy or cognitive impairment (Benincá et al., 2021; Guarani et al., 2016; Zeharia et al., 2016). To investigate the influence of Dmel_CG41128 on the life span of flies and their mitochondria, we generated flies deficient for Dmel_CG41128 using CRISPR/Cas9 genome editing (Figure 2A, B, and Supplementary Figure S1). The loss of Dmel_CG41128 reduced the average life span of flies by around 40% (Supplementary Figure S2A). Moreover, Dmel_CG41128 is present in both testes and ovaries from *D. melanogaster* (Supplementary Figure S3), and its loss reduced the overall fertility, indicated by a significantly decreased seminal vesicle area (Supplementary Figure S2B).

**Figure 2:**
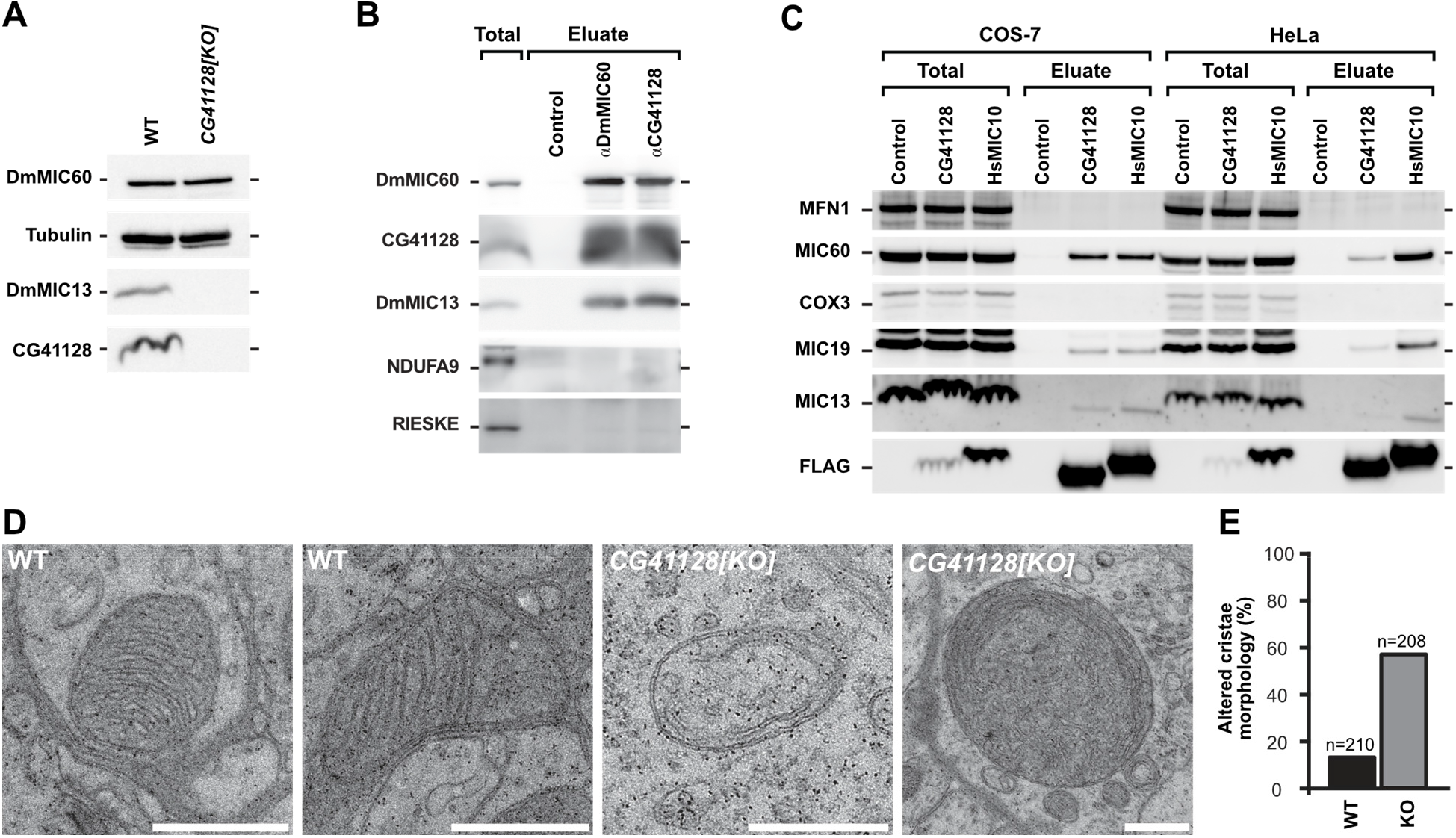
Dmel_CG41128 is DmMIC10b, a *bona fide* subunit of MICOS in *D. melanogaster*. A. Western blot analysis of steady state protein levels in solubilized mitochondria from wild type (WT) and *CG41128[KO]* flies. B. Co-IP from solubilized mitochondria from WT flies. DmMIC60 and CG41128 were used as a bait. C. Co-IP from whole cell lysates. Human MIC10-FLAG (HsMIC10) and CG41128-FLAG were transiently expressed in HeLa cells or COS-7 cells. FLAG-tagged proteins where used as a bait. D-E. Electron micrographs of mitochondria in brain tissue from WT and *CG41128[KO]* flies. n= number of analyzed mitochondrial cross sections. D. Representative images of mitochondria. E. Quantification of the cristae architecture. Scale bars: 0.5 µm.

### Dmel_CG41128 is the major MIC10 orthologue in D. melanogaster

In both humans and yeast, MIC10 controls the stability of the MIC10-subcomplex. Depletion of MIC10 leads to the degradation of MIC13 and strongly disturbs cristae architecture (Alkhaja et al., 2012; Callegari et al., 2019; Harner et al., 2011; Hoppins et al., 2011; Kondadi et al., 2020; T. Stephan et al., 2020; von der Malsburg et al., 2011). To investigate the consequences of Dmel_CG41128 depletion on mitochondrial architecture in *D. melanogaster*, we isolated mitochondria from wild-type and Dmel_CG41128-deficient flies and analyzed them by SDS-PAGE and immunoblotting. Similar to the situation in yeast and human cells (Guarani et al., 2015; Kondadi et al., 2020; von der Malsburg et al., 2011), loss of Dmel_CG41128 caused the loss of DmMIC13 when analyzing steady state levels (Figure 2A). Next, we performed co-immunoprecipitation (Co-IP) experiments on the isolated WT mitochondria using Dmel_CG41128 as a bait (Figure 2B). We found that Dmel_CG41128 interacted with both DmMIC13 and DmMIC60, central subunits of both MICOS sub-complexes (Guarani et al., 2015). Likewise, when DmMIC60 was used as a bait, Dmel_CG41128 and DmMIC13 were pulled down (Figure 2B), indicating that Dmel_CG41128 is a part of the *Drosophila* MICOS complex.

In order to test if Dmel_CG41128 also interacts with subunits of mammalian MICOS complexes, we expressed Dmel_CG41128-FLAG and FLAG-tagged human MIC10 (HsMIC10-FLAG) in human HeLa cells and COS-7 cells from the green African monkey *Cercopithecus aethiops.* Approximately the same amounts of MICOS subunits were pulled down with these proteins as baits in COS-7 cells. Also, in HeLa cells Dmel_CG41128-FLAG pulled down subunits of MICOS, although HsMIC10-FLAG was a more efficient bait (Figure 2C). We conclude that Dmel_CG41128-FLAG interacts also with the mammalian MICOS complex. MIC10-deficient mitochondria from yeast and human cells exhibit highly disturbed mitochondrial ultrastructure (Guarani et al., 2015; Harner et al., 2011; Hoppins et al., 2011; von der Malsburg et al., 2011). Transmission electron microscopy of brain tissue mitochondria of flies deficient for Dmel_CG41128 revealed similar mitochondrial phenotypes with the majority of mitochondria exhibiting aberrant cristae morphologies including tube-like and onion-shaped cristae (Figure 2D, E).

Altogether, the ubiquitously expressed Dmel_CG41128 shares sequence homology with other MIC10 proteins, presumably features a hairpin-like structure and binds to the MICOS complexes of *D. melanogaster, H. sapiens and C. aethiops*. Dmel_CG41128 further regulates the levels of DmMIC13 in flies, and its depletion strongly affects the cristae architecture. These findings support the notion that Dmel_CG41128 is the major MIC10 orthologue in *D. melanogaster*. Hence, in accordance with the uniform nomenclature for MICOS (Pfanner et al., 2014), we will refer to it as DmMIC10b from here on.

### DmMIC10b has a propensity to polymerize into filaments

When analyzing the subcellular localization of overexpressed DmMic10b in S2 fly cells (Figure 1B), it became apparent that higher expression levels of DmMic10b often seemed to alter the shape of mitochondria. To analyze this phenomenon further, we performed STED nanoscopy of S2 cells overexpressing DmMIC10b C-terminally fused with a FLAG-tag. At moderate expression levels, STED nanoscopy revealed the formation of distinct DmMIC10b clusters, comparable to the situation in yeast and human cells (Jans et al., 2013; T. Stephan et al., 2020). Intriguingly, long DmMIC10b-containing filaments, often forming bundles of filaments, were observable at higher expression levels of DmMIC10b-FLAG. These DmMIC10b-containing filaments seemed to be located inside of mitochondria and pervaded the mitochondrial network (Figure 3A, B). Formation of the filaments occurred only occasionally at lower expression levels (Figure 3B), but was regularly observed at high expression levels (Figure 3B).

**Figure 3:**
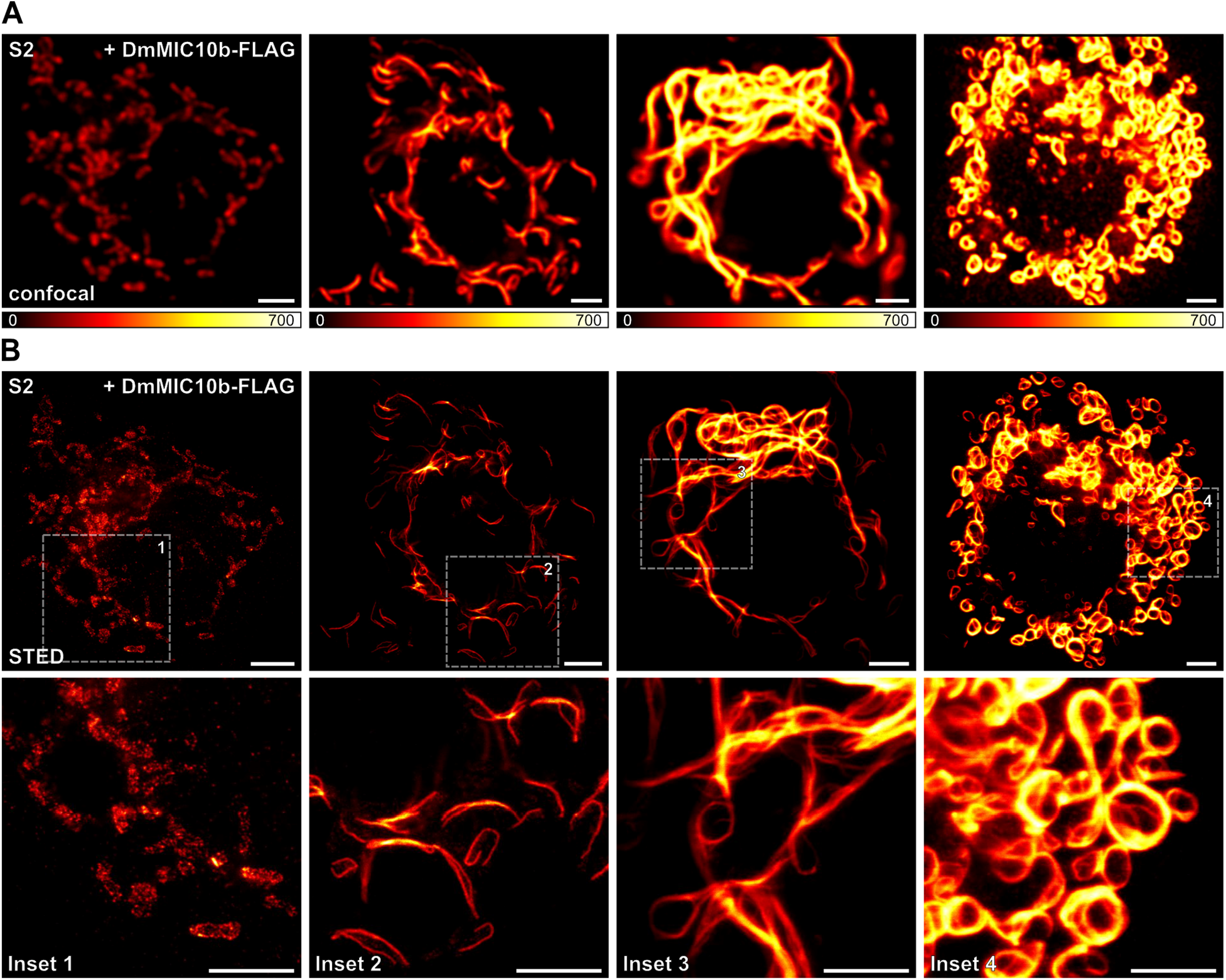
DmMIC10b can polymerize into filamentous structures. S2 cells expressing DmMIC10b-FLAG were immunolabeled against the FLAG epitope and analyzed by confocal microscopy and 2D STED nanoscopy. A. Confocal recordings of cells with different expression levels of DmMIC10b-FLAG, increasing from left to right. The signal intensity reflects the expression level. B. STED images of the cells shown in A (upper) and insets marked by dashed boxes (lower). Scale bars: 2 µM.

Even when scrutinized by STED microscopy, the filaments appeared to be contiguously labeled, suggesting that they might be formed by DmMIC10b only, rather than being a mosaic out of DmMIC10b and other proteins (Figure 3B). To explore this idea, we first expressed DmMIC10b-FLAG in two heterologous cellular systems, namely HeLa and COS-7 cells. Similar to the overexpression in S2 cells, expression of DmMIC10b in these cells resulted in the formation of DmMIC10b-containing filaments (Figure 4, Supplementary Figure S4).

**Figure 4:**
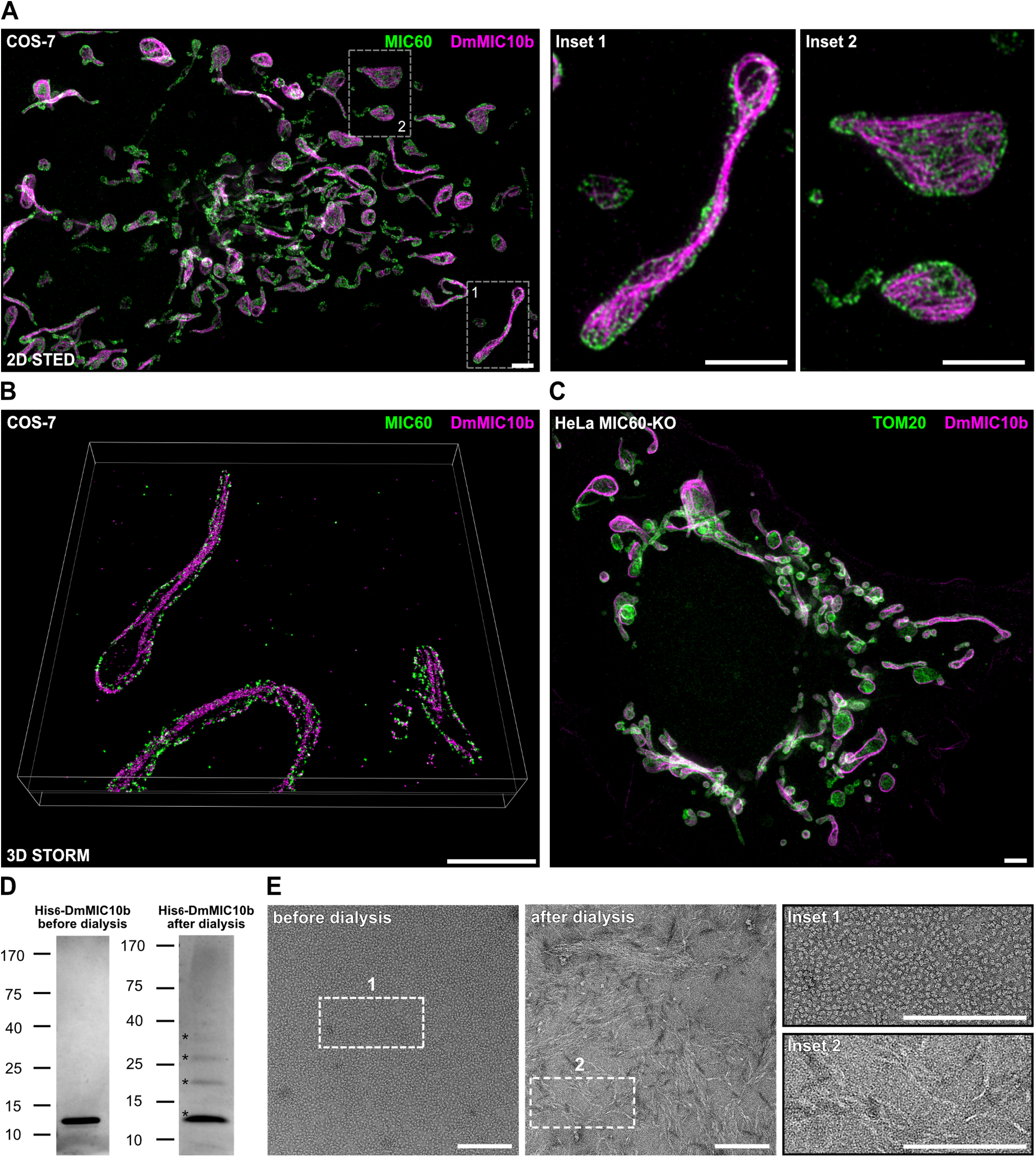
DmMIC10b can form filaments *in cellulo* and *in vitro*. A-C.) Dual-color super-resolution microscopy of COS-7 cells expressing DmMIC10b-FLAG. Cells were fixed and immunolabeled with specific antibodies against the FLAG epitope (magenta) and MIC60 (green). A. Representative 2D STED nanoscopy recording. Data were deconvolved. B. Volume rendering of a 4Pi-STORM recording. C. Representative dual-color STED recording of a MIC60-KO cell expressing DmMIC10b-FLAG. D-E. His_6_-DmMIC10b was purified from *E.coli* in the presence of urea, precipitated and solubilized using sarcosyl. The detergent was removed by dialysis. D. SDS-PAGE stained with Coomassie Brilliant Blue. Asterisks indicate different oligomeric species. E. The same sample analyzed by negative stain transmission electron microscopy. Scale bars: 2 µm (A-C); 150 nm (E).

As the Co-IP experiments showed that DmMIC10b interacts with mammalian MIC60 (Figure 2C), we also recorded the distribution of MIC60 in these DmMIC10b-overexpressing cells. Dual-color 2D STED imaging of COS-7 cells suggested occasional spatial connections between the filaments and some MIC60 clusters (Figure 4A). Nevertheless, MIC60 seemed to be localized along the IBM as described before (Harner et al., 2011; Jans et al., 2013; Pape et al., 2020), whereas the DmMIC10b-containing filaments seemed to be situated more towards the center of the mitochondrial tubules. To localize DmMIC10b with optimal accuracy in 3D, we next performed 4Pi-STORM of COS-7 cells expressing DmMIC10b-FLAG (Bates et al., 2022). The 3D recordings confirmed the initial impression that the filaments associated into bundles (Supplementary Movie S1) and that only a fraction of DmMIC10b was in close proximity to MIC60 clusters (Figure 4B, Supplementary Movie S2). Instead, a significant part of DmMIC10b-FLAG seemed to be part of filamentous structures that ran along the center of the mitochondrial tubules, suggesting that these structures were localized outside the IBM (Figure 4B, Supplementary Movie S2).

### Filament formation does not require other MICOS proteins and is independent of the C-terminal tag

The nanoscopy data supported the idea that DmMIC10b-FLAG filaments may form independent from other MICOS proteins. To test this further, we next expressed DmMIC10b-FLAG in human MIC10-KO and MIC60-KO cells, as these cells are devoid of the MIC10 sub-complex and the entire MICOS complex, respectively (T. Stephan et al., 2020). STED recordings revealed that the formation of DmMIC10b-filaments occurred in the absences of the MIC10 sub-complex as well as in the absence of the MICOS holo-complex (Figure 4C, Supplementary Figure S5A,B). We conclude that additional MICOS subunits are not required for the polymerization of DmMIC10b into filaments.

As all DmMIC10b filaments shown thus far relied on overexpression of a C-terminally FLAG-tagged version of DmMIC10b, we next expressed non-tagged DmMIC10b in mammalian HeLa, U-2 OS and COS-7 cells and labeled them with antibodies against DmMIC10b in order to test if the formation of the filamentous structures was induced by the FLAG-epitope. Also untagged DmMIC10b formed filaments at high expression levels, demonstrating that its propensity to form a filamentous structure is independent of the tag (Supplementary Figure S5C-E).

We next investigated if purified DmMIC10b is able to polymerize into filamentous structures *in vitro*. To this end, His_6_-DmMIC10b was expressed in *E. coli* and purified to homogeneity from inclusion bodies as described previously (Barbot et al., 2015). Following the removal of detergent by dialysis, purified His_6_-DmMIC10b exhibited a distinct ladder-pattern on SDS-PAGE gels (Figure 4D, right panel), as previously reported for ScMIC10 (Barbot et al., 2015; Bohnert et al., 2015). This suggests that the purified DmMIC10b can associate into stable oligomers. In line with this observation, negative stain electron microscopy demonstrated the formation of filamentous structures upon the removal of the detergent (Figure 4E). Taken together, we conclude that DmMIC10b can polymerize into homo-oligomeric filaments both in mitochondria and *in vitro*.

### Filament formation seems to be specific for DmMIC10b

Although the formation of filamentous MICOS structures had been suggested by early studies conducted on yeast (Hoppins et al., 2011), several studies using super-resolution microscopy have not substantiated the existence of extended MICOS-only filaments in yeast or mammalian cells (Jakobs, Stephan, Ilgen, & Brüser, 2020). Still, elongated MIC60 assemblies in mitochondria of COS-7 cells (Bates et al., 2022) or in mitochondria of HeLa cells depleted of the dynamin-like GTPase optic atrophy 1 (OPA1) (T. Stephan et al., 2020) have been reported. However, these ring- or arc-like structures only wrapped around the mitochondrial tubules and were by orders of magnitude shorter than the DmMIC10b filaments reported here. Previous studies overexpressing HsMIC10 also did not report on the formation of filaments at various expression levels (T. Stephan et al., 2020). To further explore if MIC10 from yeast or humans has a tendency to polymerize when expressed in a heterologous system, we expressed ScMIC10 in COS-7 cells and S2 cells and HsMIC10 in S2 cells. No filaments were observed, suggesting that the propensity to form filaments is a specific characteristic of DmMIC10b (Supplementary Fig. S6A-C).

### DmMIC10b filaments remodel the cristae membrane

As MIC10 is a membrane-shaping protein (Barbot et al., 2015; Bohnert et al., 2015), it appeared possible that the DmMIC10b filaments influence the overall IM architecture. To test this, we expressed the cristae marker COX8A fused to a SNAP-tag (Till Stephan et al., 2019) together with DmMIC10b in HeLa and COS-7 cells and visualized the fusion protein inside mitochondria using live-cell STED nanoscopy. Both mitochondria of HeLa and COS-7 control cells showed lamellar cristae as previously reported (Liu et al., 2022; Till Stephan et al., 2019). Overexpression of DmMIC10b strongly altered the cristae architecture, with the IM often apparently collapsed along the DmMIC10b filaments (Figure 5A).

**Figure 5:**
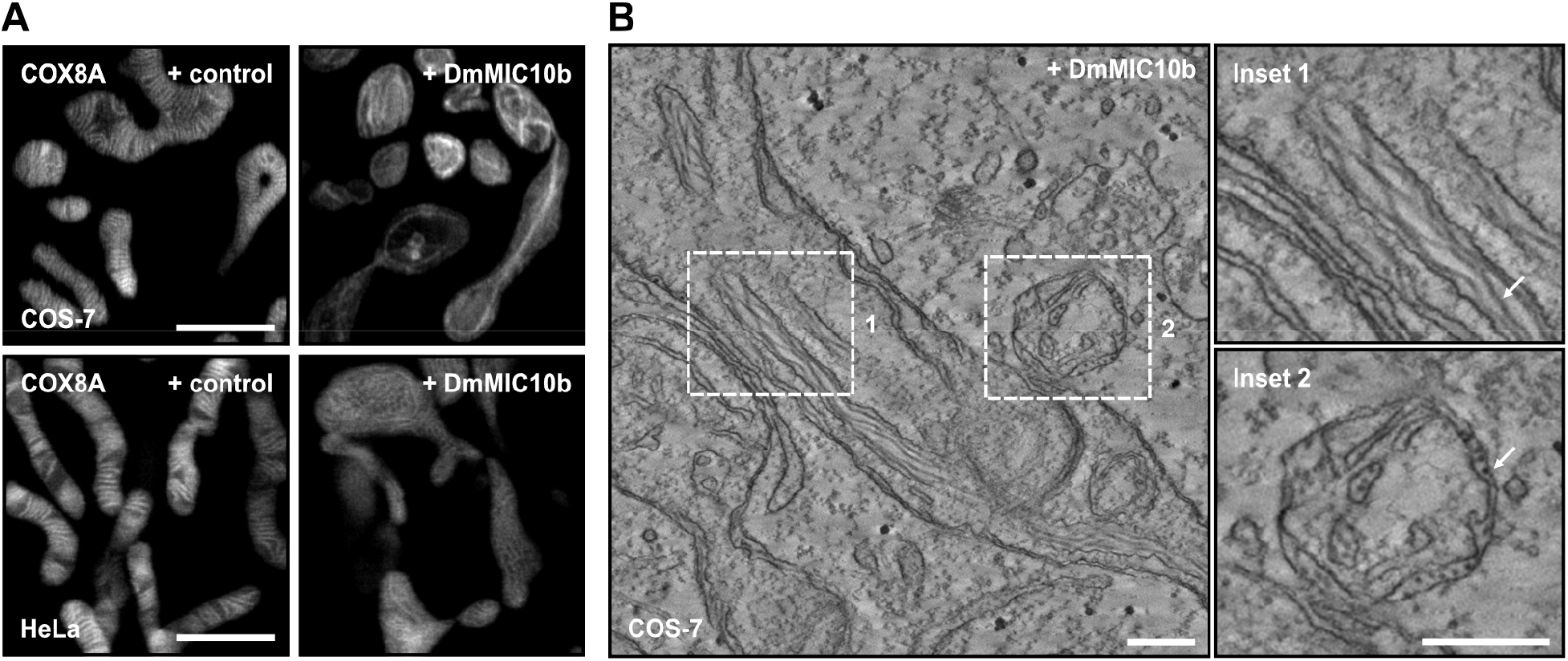
DmMIC10b polymerizes into filaments which alter the mitochondrial ultrastructure. A. Live-cell STED of mammalian cells expressing DmMIC10b. COS-7 and HeLa cells were co-transfected with DmMIC10b-FLAG and COX8A-SNAP to visualize the cristae membrane. Cells were labeled with SNAP-cell SiR 647 and visualized by 2D live-cell STED nanoscopy. B. Electron tomography of mitochondria from COS-7 cells expressing DmMIC10b-FLAG. Arrows point to DmMIC10b-FLAG filaments located within the cristae lumen (upper) and the IMS (lower). Scale bars: 2 µm (A); 0.25 µm (B).

In order to investigate the influence of DmMIC10b polymerization on the cristae architecture in more detail, we next recorded electron tomograms of chemically fixed COS-7 cells expressing DmMIC10b. Electron microscopy revealed filament bundles oriented along the mitochondria. As these filaments were absent in WT cells, we assume that these are DmMIC10b filaments. Whereas some of these filaments were in close contact with the IM, other filaments seemed to form between OM and IBM and along the cristae lumen, thereby widening the IMS or causing the formation of aberrant, tubular cristae, respectively (Figure 5B). As STED nanoscopy recordings of MICOS knockout cells expressing DmMIC10b had shown a similar arrangement of the DmMIC10b filaments (Supplementary Figure S5A, B), we analyzed those cells using electron microscopy as well. Remarkably, these cells featured a similar internal organization of the DmMIC10b filaments despite the strikingly different cristae architecture of WT, MIC10-KO and MIC60-KO cells (Kondadi et al., 2020; T. Stephan et al., 2020). In MIC60-KO cells, we observed the majority of filaments in bundles between OM and IBM, which we attribute to the strongly reduced number of CJs in these mitochondria (Supplementary Figure S7A, B).

Together, our data suggest that upon overexpression DmMIC10b polymerizes into extended filaments that associate into bundles within the IMS and influence the mitochondrial IM architecture.

### Filament formation of MIC10b is regulated by conserved amino acids

#### DmMic10b oligomerization is a prerequisite for filament formation

Filament formation seemed to be a peculiarity of DmMIC10b, hence we compared its primary sequence to that of other MIC10 proteins (Figure 6A). The MIC10 proteins of *D. melanogaster, H. sapiens, R. norvegicus, D. rerio and S. cerevisiae* exhibit highly conserved glycine-rich motifs in each of the two TM segments that mediate MIC10 oligomer formation in yeast (Barbot et al., 2015; Bohnert et al., 2015). The first TMD contains a GxxxG motif, whereas the second TMD contains a more extended GxGxGxG motif (Figure 6A). The yeast ScMIC10 somewhat stands out as it exhibits an additional amino acid sequence of approximately 20 residues at its N-terminus (Figure 6A). Furthermore, all analyzed MIC10 proteins except ScMIC10 feature two highly conserved cysteine residues: a conserved cysteine in a distance of 11 aa N-terminally of the GxxxG motif (C19 in DmMIC10b) and a conserved cysteine that is placed 6 aa C-terminally of the GxGxGxG motif (C64 in DmMIC10b). DmMIC10b uniquely features a cysteine residue (C28) in close proximity to the N-terminal GxxxG motif, whereas in HsMIC10, RnMIC10 and DrMIC10 the corresponding amino acid is glycine and extends the GxxxG motif into a GxGxGxG motif. In addition, the other MIC10 proteins, except for DmMIC10b, feature a phenylalanine at position 41, whereas DmMIC10b has an isoleucine at this position (Figure 6A).

**Figure 6:**
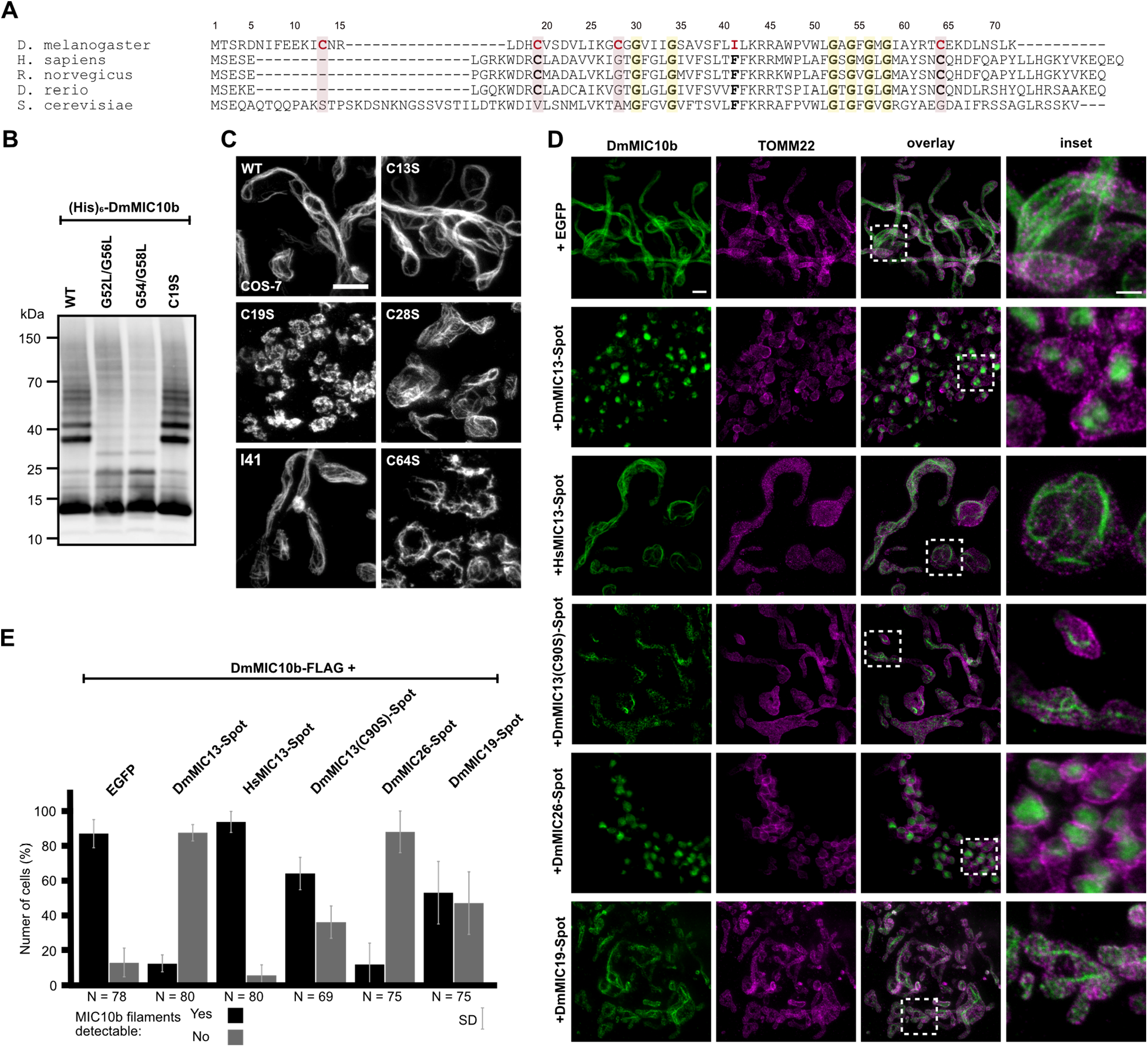
Formation of DmMIC10b filaments depends on conserved amino acids and is suppressed by DmMICOS proteins. A. Amino acid sequence alignment of MIC10-like proteins from *D. melanogaster* and MIC10 from humans, rats, zebra-fish and yeast. Amino acid residues exchanged for analysis in (B-C) are marked in red. B. Western blot analysis of DmMIC10b mutants. His_6_-DmMIC10b with point mutations was expressed in *E. coli*. Cells were homogenized and the insoluble pellet was solubilized using sarcosyl. The detergent was removed by dialysis and the samples were analyzed by SDS-PAGE and immunoblotting. C. 2D STED nanoscopy of COS-7 cells expressing mutants of DmMIC10b-FLAG. Cells were chemically fixed and immunolabeled for the FLAG epitope. D.-E. Co-expression of MICOS proteins in COS-7 cells. Cells were co-transfected to induce expression of DmMIC10b-FLAG together with other Spot-tagged MICOS proteins. Cells were immunolabeled against FLAG, Spot and TOMM22. Double-transfected cells were recorded by dual-color STED nanoscopy. D. Representative STED nanoscopy recordings. E. Quantification of the number of cells which formed DmMIC10b-FLAG filaments. Bars indicate the mean of three independent biological repeats, whiskers indicate the standard deviation. N indicates the total number of analyzed cells. Scale bars: 2 µm (B), 1 µm (C, overview), 0.5 µm (C, inset).

As the glycine-rich motifs mediate MIC10 oligomer formation in yeast (Barbot et al., 2015; Bohnert et al., 2015), we next explored if disruption of the GxGxGxG motif in *D. melanogaster* has an influence on filament formation. To this end, mutations to disturb this motif in the second TMD were introduced. Concretely, we expressed DmMIC10b(G52L/G56L) and DmMIC10b(G54L/G58L) in S2 cells and in *E.coli* and analyzed their ability to form higher molecular weight species or filaments. After expression of the proteins in *E. coli* cells, we lysed the cells, enriched inclusion bodies and solubilized them with sarcosyl. Following dialysis, we analyzed the samples using SDS-PAGE and western blotting (Figure 6B). As previously reported for yeast MIC10, immunoblotting showed that disruption of the GxGxGxG motif in DmMIC10b strongly reduced the amount of higher molecular weight species preserved during SDS-PAGE (Barbot et al., 2015; Bohnert et al., 2015). Likewise, both mutations abolished filament formation, but instead resulted in the appearance of protein clusters along the mitochondria (Supplementary Figure S8A).

Together this suggests that oligomerization through its GxGxGxG motif located in the second TMD is a prerequisite for the polymerization of DmMIC10b into filaments.

#### Filament formation but not oligomer formation requires a conserved cysteine residue

Since DmMIC10b differs significantly from the well-studied ScMIC10 with respect to residues C13, C19, C28, C64, and I41, we investigated whether mutation at these sites affects the ability of DmMIC10b to polymerize into filaments. The cysteine residues were individually replaced by a serine residue and the isoleucine by a phenylalanine residue. The variants were expressed in COS-7 and HeLa cells and their localization recorded by STED nanoscopy (Figure 6B). Except DmMIC10b(C19S), which resulted in a clustered, non-filamentous protein distribution, the other tested DmMIC10b variants were all capable of filament formation (Figure 6B, Supplementary Figure S8B). Therefore, we next tested if C19, like the GxGxGxG motif, is essential for the formation of MIC10 oligomers. In strong contrast to variants mutated in the GxGxGxG motif, DmMIC10b(C19S) behaved virtually identical to WT DmMIC10b upon SDS-PAGE (Figure 6B).

We conclude that DmMIC10b oligomerizes into higher molecular weight species through its glycine-rich motif as previously reported for MIC10 from yeast. DmMIC10b oligomers in turn require the conserved C19 residue to assemble into filaments.

### DmMICOS subunits can suppress filament formation of DmMIC10b

The formation of extended DmMIC10b filaments in fly cells was only observed upon overexpression of DmMIC10b, which is a condition that results in a shifted balance between DmMIC10b and the other MICOS proteins. We speculated that at physiological conditions filament formation of DmMIC10b is suppressed by interacting MICOS proteins. To explore this, DmMIC10b-FLAG was co-expressed with several Spot-tagged versions of MICOS proteins of the MIC10 and the MIC60 sub-complexes, and filament formation was determined by STED nanoscopy (Figure 6D). To ensure balanced expression of the respective two overexpressed proteins, we expressed both proteins as a translational fusion, separated by a self-cleaving 2A peptide.

Co-expression of DmMIC10b with the MIC10-subcomplex subunits DmMIC13 or DmMIC26 suppressed DmMIC10b filament formation in the vast majority (> 80%) of the cells, whereas DmMIC19, part of the MIC60-subcomplex, reduced filament formation less efficiently (∼50%). Upon co-expression of the human DmMIC13 orthologue HsMIC13, the majority of the analyzed cells still formed filaments, suggesting that the fly homologs have co-evolved to efficiently suppress the ability of DmMIC10b to form filaments at physiological conditions (Figure 6D, E). The suppression of filaments by DmMIC13 was reduced when its only cysteine residue (C90) was replaced by a serine residue (Figure 6D, E), further supporting the notion that in *D. melanogaster* cysteine residues are key for the regulation of the oligomerization status of MIC10.

Taken together, DmMIC10b, the major MIC10 from *D. melanogaster*, has the propensity to form extended filaments in *vitro* and *in cellulo*. Upon overexpression, the filaments reside in the IMS and deform the IM. Formation of filaments requires glycine-rich motifs within the C-terminal TMD as well as a conserved cysteine residue (C19) in close proximity to the N-terminal TMD. The formation of filaments requires excess of DmMIC10b and is effectively suppressed by co-overexpression of other constituents of the MIC10-subcomplex.

## Discussion

In this study we show that the mitochondrial protein Dmel_CG41128/ DmMIC10b is the major MIC10 orthologue in *D. melanogaster*. DmMIC10b is ubiquitously expressed and interacts with the MICOS complexes of flies, humans and monkeys. It controls the stability of the MIC10-subcomplex of *D. melanogaster* and is required for maintaining the mitochondrial ultrastructure. Despite its high homology to ScMIC10 and HsMIC10, DmMIC10b stands out due to its propensity to polymerize into extended filaments. At low expression levels, DmMIC10b formed distinct clusters, resembling the distribution of MICOS proteins in yeast or human cells (Bates et al., 2022; Jans et al., 2013; Kondadi et al., 2020; Pape et al., 2020; Stoldt et al., 2019), whereas at high expression levels, DmMIC10b polymerized into bundles of filaments which influenced both the fusion-fission balance of the mitochondrial network and the cristae ultrastructure. Remarkably, these filaments were located between the OM and IM as well as inside the cristae lumen, suggesting that they can form or extend outside of the IM.

Filaments formed by DmMIC10b resemble those formed by the GFP-tagged bacterial cytoskeleton protein Mreb (Grotjohann et al., 2011; Pande, Mitra, Bagde, Srinivasan, & Gayathri, 2022). The formation of extended Mreb filaments has been described as an artifact caused by tagging with fluorescent proteins (Swulius & Jensen, 2012). Under physiological conditions, untagged Mreb polymerizes into smaller arc-like or ring-like assemblies, which are crucial to determine the bacterial cell shape (Shi, Bratton, Gitai, & Huang, 2018). This study shows that the formation of DmMIC10b filaments is independent of the C-terminal tag. Instead, polymerization into filaments depends primarily on the expression level and thereby concentration of DmMIC10b.

Similar to yeast MIC10 (Bohnert et al., 2015), DmMIC10b relies on the conserved GxGxGxG motif in the C-terminal TM segment to form oligomeric species that are preserved during SDS-PAGE. In the case of DmMIC10b, this oligomerization seems to be required also for the assembly of extended filaments. In addition, we found that a highly conserved cysteine residue (C19 in DmMIC10b), located N-terminally of the first TMD of DmMIC10b is crucial for the formation of filaments, but not of oligomers. Remarkably, this cysteine residue, as well as the cysteine residue located C-terminally of the second TM segment (C64 in DmMIC10b), is highly conserved in the MIC10 proteins of higher animals, including humans, flies, rats, and fish, whereas the well-studied MIC10 from yeast does not contain any cysteine residue (Figure 6A). This may point to understudied differences between the MICOS complexes of lower and higher eukaryotes. Given the sequence homology of the MIC10 proteins in higher eukaryotes, the question remains why DmMIC10b has the propensity to form filaments, whereas the human MIC10 seems not to have this ability. A crystal structure of MIC10 is not available, but models generated by AlphaFold2 may provide some hints (Fig. 1C). Structure predictions suggest that DmMIC10b, HsMIC10 and ScMIC10 all feature a hairpin-like topology, but differences exist regarding the length and shape of the two alpha helices that contain the TMDs and the conserved glycine-rich motifs. The predictions suggest that in DmMIC10b, the N-terminal alpha helix, which contains the conserved C19 residue, is elongated and exhibits a curved shape compared to other MIC10 proteins. Moreover, DmMIC10b does not seem to feature the flexible termini predicted for human or yeast MIC10. We speculate that such differences in the shape of the MIC10 monomers influence the shape of MIC10 oligomers and their ability to polymerize into filaments.

We found that in fly cells, the stoichiometric ratio between DmMIC10b and other MICOS subunits influences the propensity of DmMIC10b to polymerize into extended filaments. This seems to be an effective mechanism to keep MIC10 oligomerization in check. Indeed, we never observed extended DmMIC10b filaments in wild-type fly cells, which does not exclude that possibility that at specific developmental stages the ratio between DmMIC10b and the other MICOS subunits is changed and filaments are formed in order to rearrange the mitochondrial architecture. Therefore, it will be revealing to investigate the interplay of cristae-shaping proteins along all developmental stages of an organism such as a fly.

Movie S1. 4Pi-STORM of COS-7 cells expressing DmMIC10b-FLAG. Cells were immunolabeled with specific antibodies against the FLAG epitope and recorded by 4Pi-STORM. The movie shows a volume rendering of the 4Pi-STORM recording. The position along the z-axis is color-coded.

Movie S2. Dual-color super-resolution microscopy of COS-7 cells expressing DmMIC10b-FLAG. Cells were immunolabeled with specific antibodies against the FLAG epitope (magenta) and MIC60 (green) and recorded with 4Pi-STORM. The movie shows a volume rendering of the 4Pi-STORM recording.

## Acknowledgements

We thank Sylvia Loebermann, Rita Schmitz-Salue, Karin Hartwig and Doris Brentrup for excellent technical assistance and maintenance of fly stocks. This work was supported by the European Research Council (ERCAdG No. 835102, to SJ). It was funded by the DFG-funded FOR2848 (project P05 to MM and Z01, to DR) and SFB1190 (project P01 to SJ and P13 to PR). Stocks obtained from the Bloomington Drosophila Stock Center (NIH P40OD018537) were used in this study.

## Author contributions

T.S., S.S., M.Barbot, and S.J. conceived the project. T.S., S.S., M.Barbot, M.Bates, T.C., S.D., P.R., and S.J. designed research. T.S., S.S., M.Barbot, T.C., F.L., M.Bates, P.B.D., D.R. and S.D. performed experiments. T.S., S.S., M.Barbot, T.C., F.L., M.Bates, P.B.D., H.S., M.M., D.R., S.D., P.R., and S.J. analyzed data. T.S. and S.J. wrote the paper with comments from all authors.

---

**Supplementary Figure S1:**
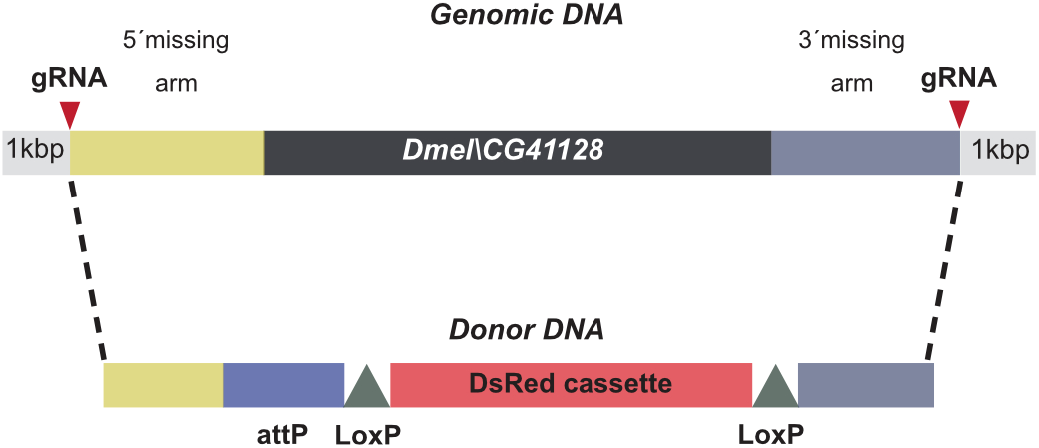
Generation of a *CG41128* knockout in flies. CRISPR/Cas9 gene editing was used to insert a DsRed cassette into the *CG4118* gene.

**Supplementary Figure S2:**
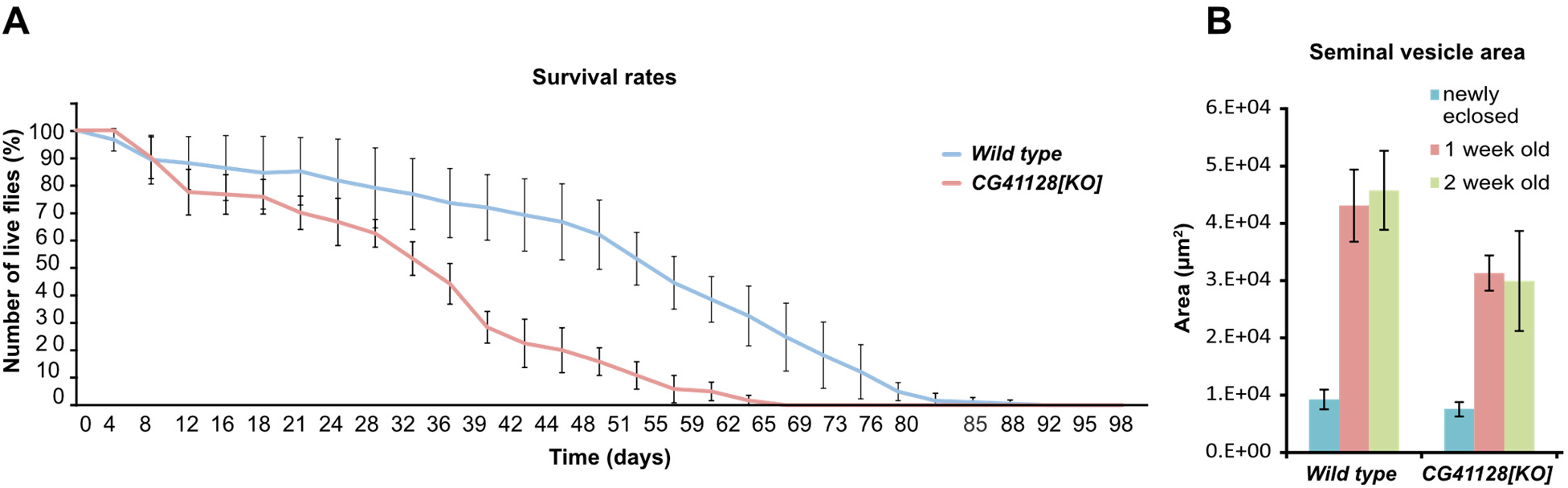
Loss of Dmel_CG41128 reduces the life span and fertility of flies. A. Survival rates of wild-type flies and CG41128-deficient flies. Curves indicate the mean of four independent biological repeats (n=4). Whiskers indicate the standard deviation. B. Seminal vesicle area in wild-type and *CG41128[KO]* flies. The seminal vesicle area was estimated for newly eclosed flies, one- and two-week old flies. Bars indicate seminal vesicle area of 15-25 seminal vesicles per time point and genotype, and whiskers indicate the standard deviation.

**Supplementary Figure S3:**
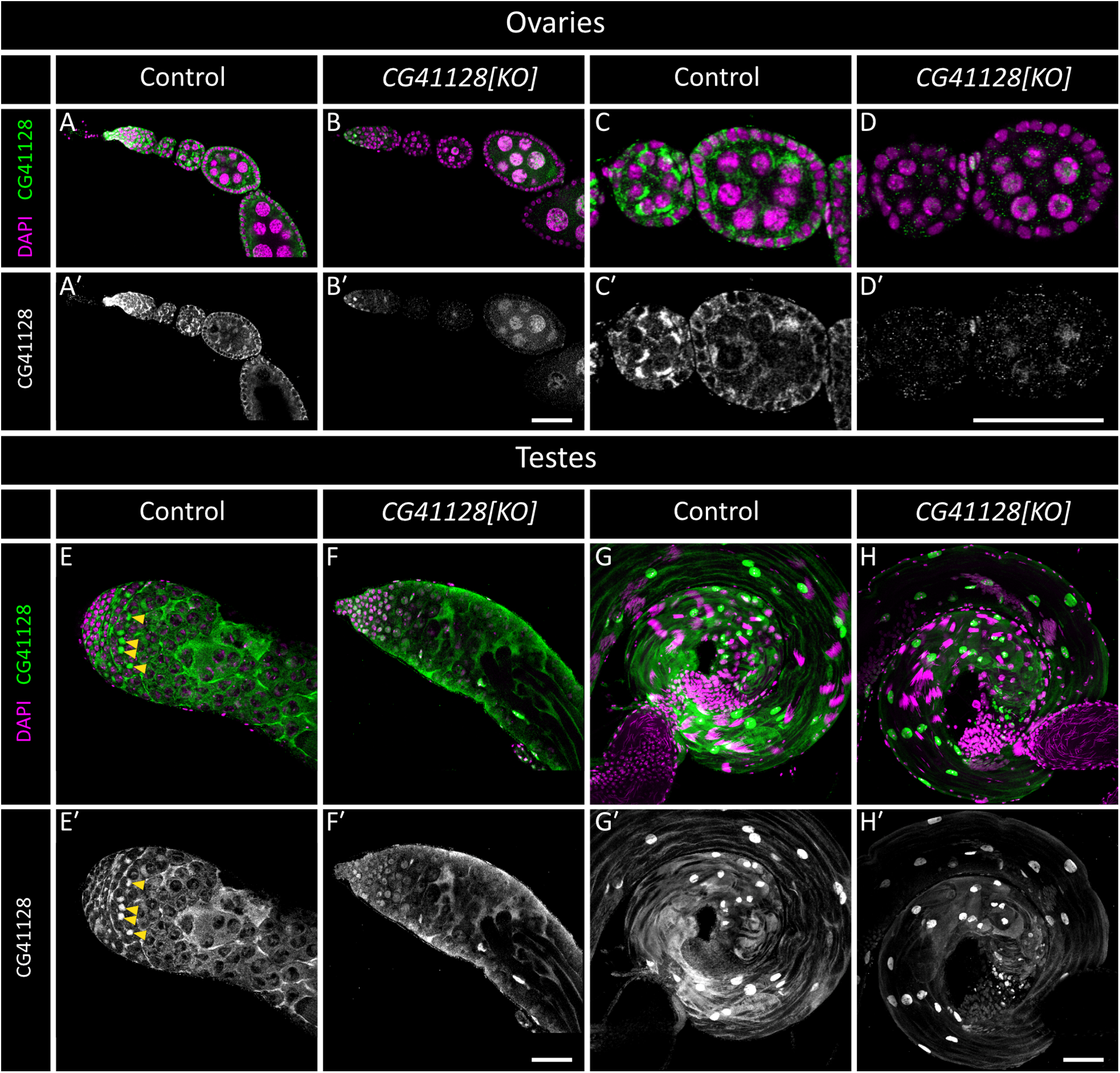
Dmel_CG41128 is present in ovaries and testes of *D. melanogaster*. Ovaries (A-D) and testes (E-H) from control (*w[1118]/OR-R*) and mutant (*CG41128[KO]*) adult flies. Shown are ovarioles (A-B) and developing egg chambers (C-D); and apical ends (E-F) and basal ends (G-H) of testes. DAPI (magenta) marks the nuclei, and anti-CG41128 staining is shown in green. Single grayscale panels (A′-H′) depict anti-CG41128 signal. CG41128 is present in mitochondria in both ovaries (adjacent to nuclei in C, C′) and testes (yellow arrowheads in E, E′). Mitochondrial staining is abrogated in *CG41128[KO]* tissues, but some nonspecific staining remains, including strong staining in testis cyst cell nuclei (see staining in H′). Scale bars 50 µm.

**Supplementary Figure S4:**
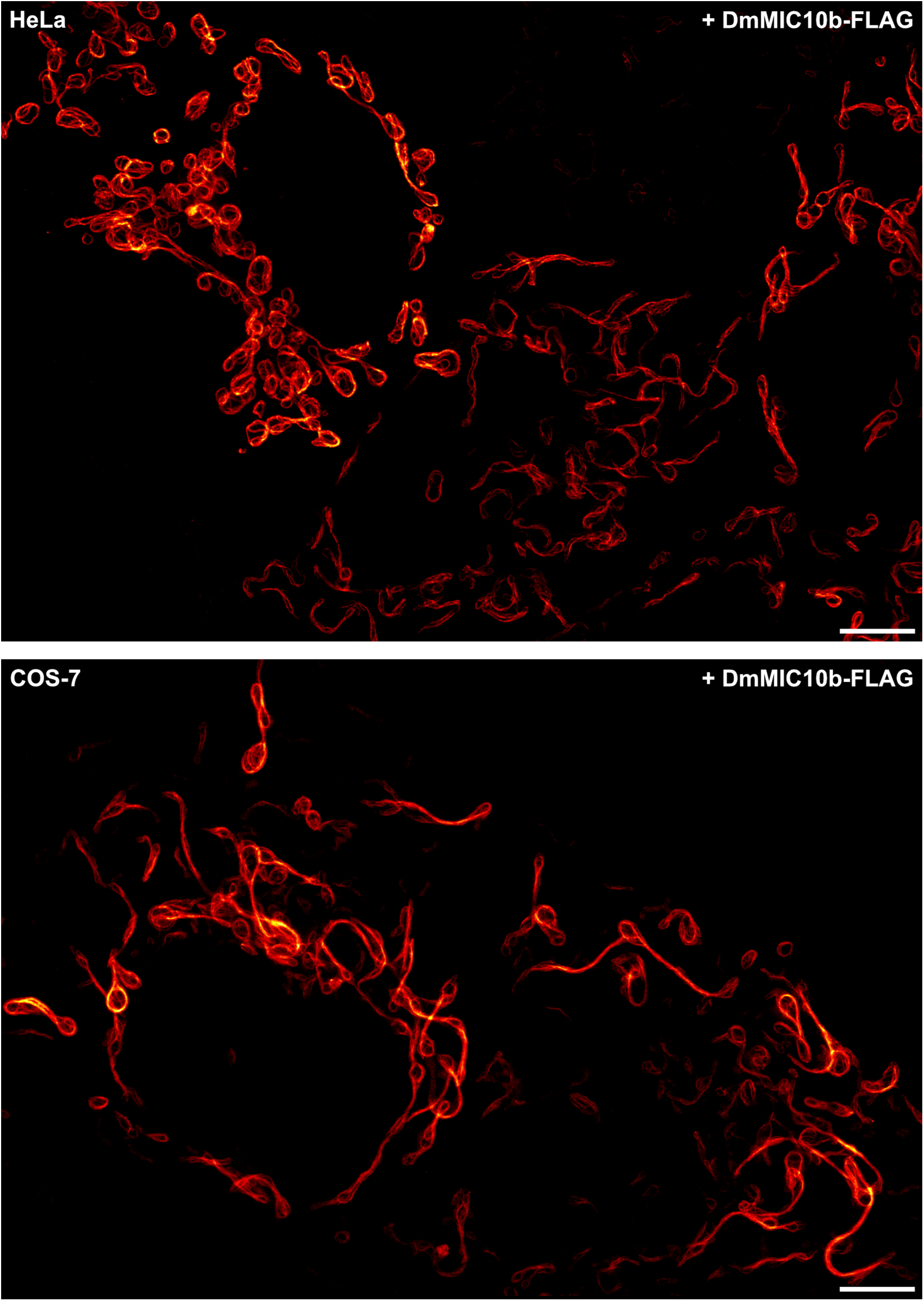
DmMIC10b overexpression causes formation of filaments. DmMIC10b was expressed in HeLa (upper) and COS-7 (lower) cells. Cells were chemically fixed, immunolabeled and visualized by 2D STED nanoscopy. Scale bars: 5µm.

**Supplementary Figure S5:**
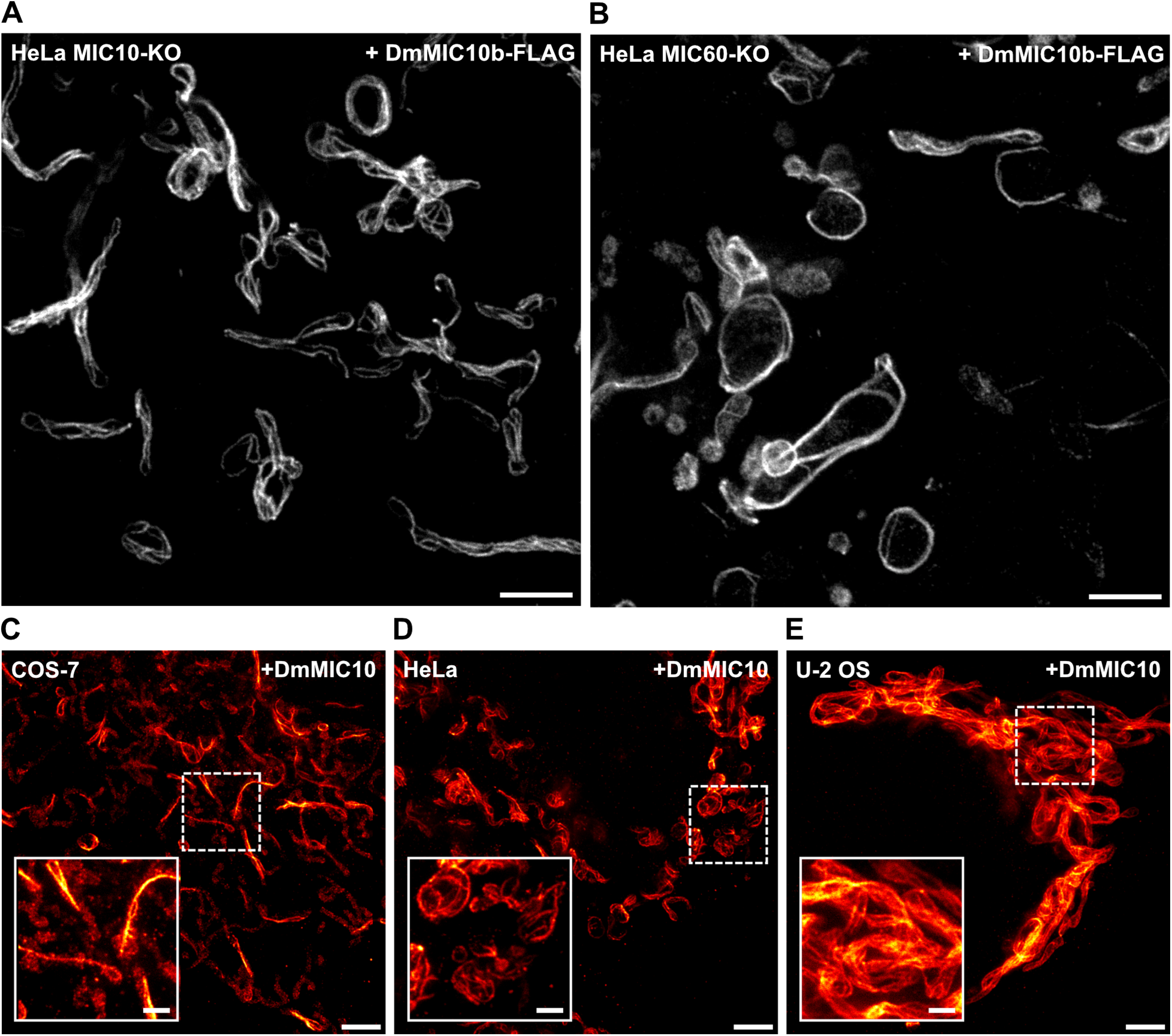
DmMIC10b expression induces filament formation in the absence of human MICOS proteins and irrespective of the C-terminal FLAG-tag. A.-B. Genome-edited HeLa cells expressing DmMIC10b-FLAG were immunolabeled against the FLAG epitope and recorded with 2D STED nanoscopy. A. HeLa cells depleted of HsMIC10 (MIC10-KO) which are lacking the MIC10-subcomplex. B. HeLa cells depleted of HsMIC60 (MIC60-KO) which are lacking virtually all HsMICOS components. C. –E. 2D STED nanoscopy of immunolabeled COS-7 (C), HeLa (D) and U-2 OS (E) cells expressing non-tagged DmMIC10b. Scale bars: 2 µm (A-B); 2 µm (overview, C-E), 0.5 µm (inset, C-E).

**Supplementary Figure S6:**
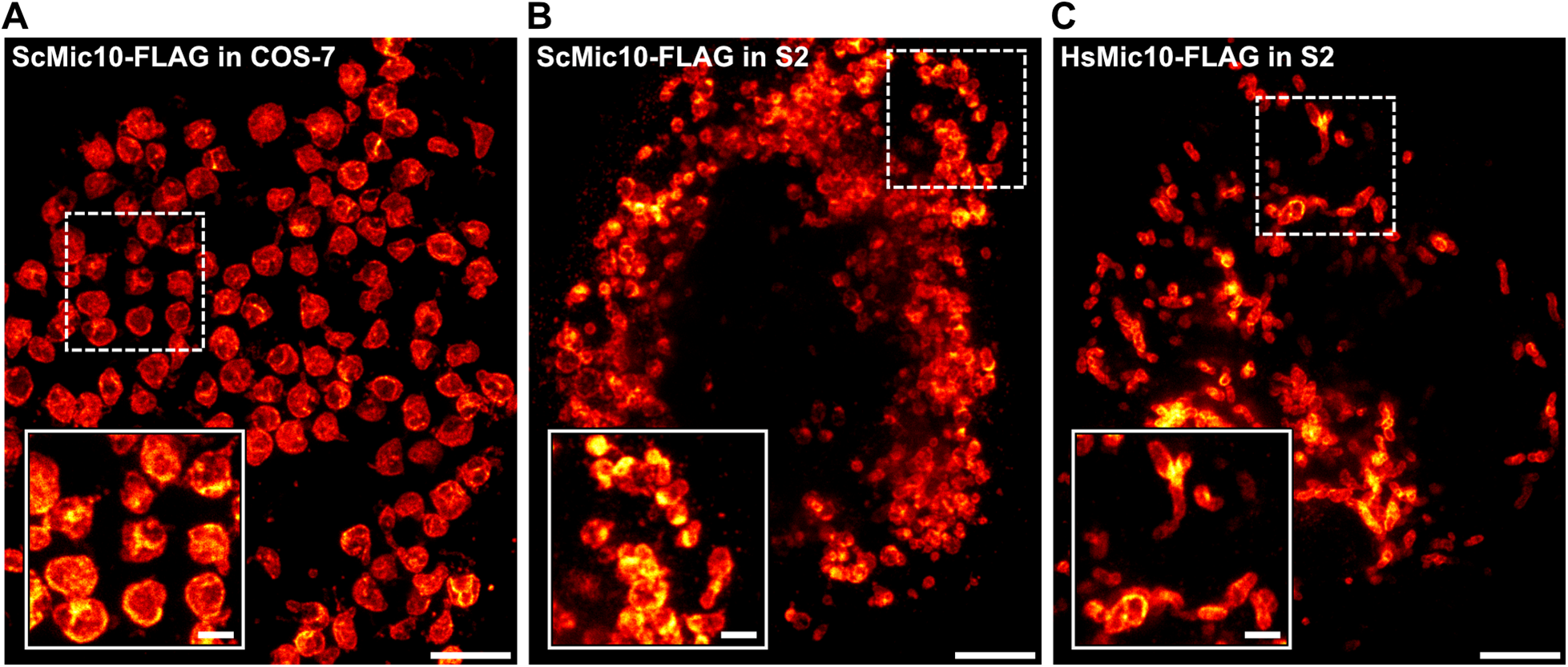
MIC10-FLAG from yeast or humans does not polymerize into filaments when expressed in cultivated cells. A. –C. Cells were transfected, chemically fixed and immunolabeled against the FLAG epitope. The cells were analyzed by 2D STED nanoscopy. A. COS-7 cell expressing ScMIC10-FLAG. B. S2 cell expressing ScMIC10. C. S2 cell expressing HsMIC10. Scale bars: overview, 2 µm; inset, 0.5 µm.

**Supplementary Figure S7:**
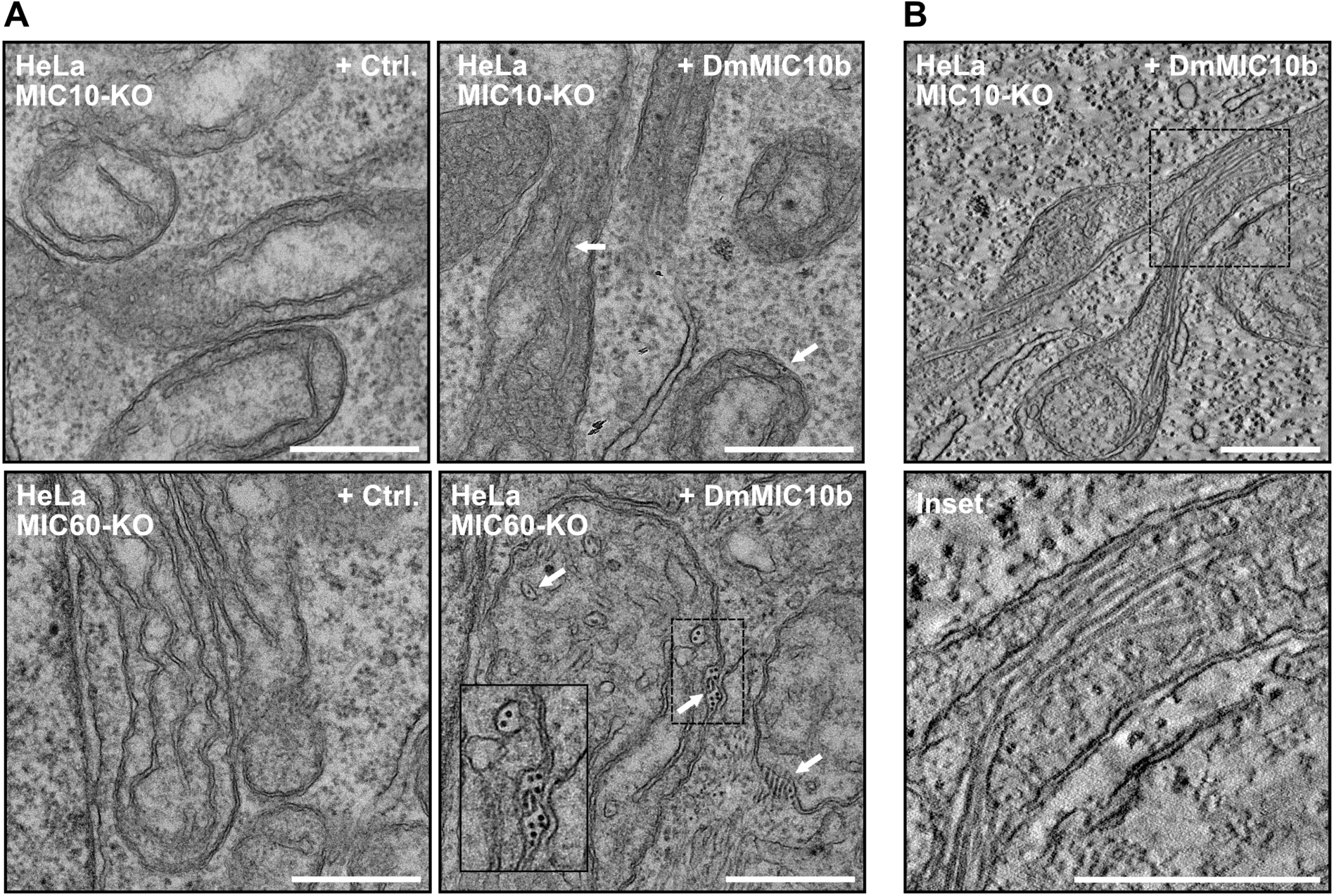
DmMIC10b filaments influence the cristae architecture in MICOS-deficient mitochondria. A. –B. HeLa cells deficient for MIC10 (MIC10-KO) or MIC60 (MIC60-KO) were transfected to induce the expression of DmMIC10b-FLAG. Cells were chemically fixed and analyzed by transmission EM. A. Electron micrograph of ultra-thin sections. White arrows indicate filaments. B. Single plane of an electron tomography data set. Scale bars: 0.5 µm.

**Supplementary Figure S8:**
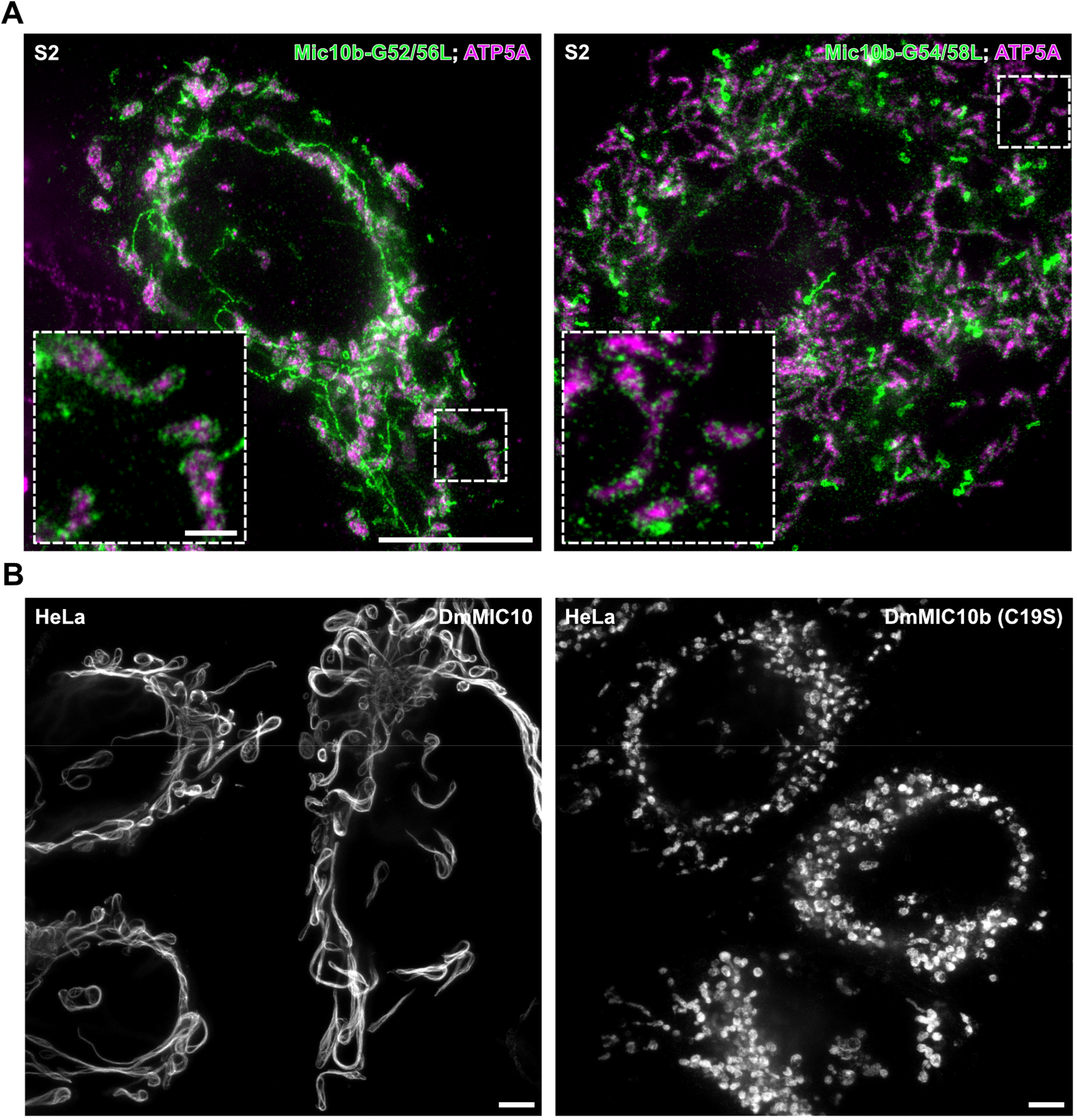
DmMIC10b polymerization requires conserved amino acids and can be suppressed by other DmMICOS proteins. A. 2D STED nanoscopy of S2 cells expressing glycine mutants of DmMIC10b-FLAG. Cells were transfected for overexpression of DmMIC10b(G52L/G56L)-FLAG (left) or DmMIC10b(G54L/G58L)-FLAG (right). Cells were immunolabeled against the FLAG-tag and ATP5A and visualized by two-color STED nanoscopy. Insets show a magnified view of the areas marked by the dashed boxed. B. 2D STED nanoscopy of HeLa cells expressing DmMIC10b-FLAG (left) or DmMIC10b(C19S)-FLAG (right). Cells were transfected for transient expression of DmMIC10b and immunolabeled against the FLAG epitope. Scale bars: 5 µm (A, overview), 0.5 µm (A, inset); 5 µm (B).

